# eIF5A is an indispensable protein for eukaryotic cells

**DOI:** 10.64898/2026.07.17.739231

**Authors:** Samoa Prieto-Díez, Alejandro Aguilar-Díaz, Luis Araque, David Peris, Paula Alepuz

## Abstract

eIF5A is an evolutionarily conserved protein found in all eukaryotes, and in Archaea and bacteria. It functions as a translation factor promoting ribosomal elongation, and it can also bind to genes in the nucleus and to mRNA. eIF5A is encoded by two paralogous genes in most eukaryotes, with one copy (*TIF51A* in yeast and *EIF5A1* in humans) highly expressed and essential, and the other (*TIF51B* in yeast and *EIF5A2* in humans) repressed in most cells and conditions. eIF5A is linked to viral infection, aging, diabetes and neurodevelopmental disorders. Whilst derepression of the duplicated silent gene *EIF5A2* is associated with several cancers, promoting metastasis. To expand our knowledge of eIF5A’s function, we searched for suppressors of yeast temperature-sensitive mutants of *TIF51A*, which cannot grow at restrictive temperature. All suppressors contained mutations in the transcriptional repressors Rox1 and Mot3, resulting in the upregulation of the paralogous gene *TIF51B*. Next, we searched for suppressors of *TIF51A* temperature-sensitive mutants in yeast lacking the *TIF51B* gene. The frequency of suppression was 20-times lower and all suppressors were revertants or contained intragenic mutations in the Tif51A protein that conferred stability. Our results suggest that the duplicated eIF5A gene serves as a non-conventional backup system that rescues mutations in the first copy, but acting at the population level. Furthermore, our results expand our understanding of the repression mechanisms that keep the second eIF5A gene silent. Lastly, our results demonstrate that eIF5A is an indispensable protein in eukaryotic cells, whose function cannot be substituted by mutations in other proteins or pathways.

**Article summary:** eIF5A is an evolutionary conserved protein encoded by two paralogous genes in eukaryotes: one highly expressed and essential, and the other silent. eIF5A promotes translation elongation, and its deficiency is linked to diabetes, aging and neurodevelopmental disorders; while derepression of the silent gene promotes metastatic cancers. We isolated suppressors of conditional mutants of the first eIF5A gene in yeast cells. Suppressors either up-regulated the second eIF5A gene or reverted first copy mutations, restoring eIF5A protein levels and survival. Our work deepens our understanding of the eIF5A paralogue gene repression, and demonstrates that eIF5A is an indispensable protein for eukaryotic cells.

## INTRODUCTION

Interest in the protein eIF5A has grown in the last decade, as it is involved in several cellular processes, and deficiencies or over-expression of the protein are associated with many diseases. eIF5A’s main role is assumed to be the promotion of translation. To bind the ribosomes, eIF5A must be activated by hypusination, a post-translational modification that takes place in two enzymatic steps and consist in the addition of one spermidine to a conserved lysine residue (Schnier et al. 1991; Park and Wolff 2018). Remarkably, the hypusination pathway is highly conserved and the enzymes essential in eukaryotes, although eIF5A is the only known target of the pathway. Hypusinated eIF5A (hyp-eIF5A) binds to ribosomes that stall during the translation of consecutive codons encoding for amino acids problematic for peptide bond formation. Binding of hyp-eIF5A induces a conformational change in the ribosome that promotes the formation of the peptide bound and restores translation (Gutierrez et al. 2013; Dever et al. 2014; Pelechano and Alepuz 2017; Schuller et al. 2017; and reviewed in Dever et al. 2018 and Barba-Aliaga and Alepuz 2022a). Thus, the role of eIF5A promoting the synthesis of specific proteins, such as formins, collagen, Atg3, TEFB and Tim50, links the protein to cellular processes as cytoskeleton organization, polarized growth, autophagy and mitochondrial function, among others (Li et al. 2014; Muñoz-Soriano et al. 2017; Lubas et al. 2018; Zhang et al. 2019; Barba-Aliaga et al. 2021; Barba-Aliaga et al. 2024). Recently, it has also been shown that eIF5A participates in the ribosome quality control (RQC) pathway by binding to dissociated 60S subunits of stalled ribosomes, where it promotes the degradation of truncated nascent peptides (Tesina et al. 2023). In addition to its direct role in cytoplasmic translation, the eIF5A protein is also found in the nucleus and can bind to mRNA (Rosorius et al. 1999; Ishfaq et al. 2012; Aksu et al. 2019; Choi et al. 2024). Moreover, we have recently shown that nuclear eIF5A binds to its translational target genes to regulate their transcription (Barba-Aliaga et al. 2026).

eIF5A is also a relevant example of a protein encoded by duplicated gene pairs that have been kept through evolution. Most eukaryotes contain two genes encoding this factor, *EIF5A1* and *EIF5A2* in humans (Jenkins et al. 2001) and *TIF51A* (*HYP2*) and *TIF51B* (*ANB1*) in *Saccharomyces cerevisiae* (Kang et al. 1992). In most eukaryotes, the two eIF5A paralogue genes maintain a high identity in their amino acid sequences (84% in humans and 90% in *S. cerevisiae*) and the eIF5A isoforms seem to be interchangeable (Schnier et al. 1991; Schwelberger et al. 1993; Jenkins et al. 2001). However, the paralogue genes show different expression patterns in both organisms. Thus, in humans *EIF5A1* is highly expressed in most cell types whereas *EIF5A2* is mostly silent, but expressed in testis and neurons (Jenkins et al. 2001). In yeast, *TIF51A* is also highly expressed under most conditions whereas *TIF51B* is repressed; but switching to oxygen restrictive conditions down-regulates *TIF51A* and up-regulates *TIF51B* (Schwelberger et al. 1993; Barba-Aliaga et al. 2020; Barba-Aliaga and Alepuz 2022b). As mentioned above, the growing interest in eIF5A comes from its relation with diverse pathologies. Thus, deficient levels of active eIF5A has been related to several diseases such as diabetes, neurodevelopmental syndromes, and also to aging and viral infection (Ruhl et al. 1993; Maier et al. 2010; Tersey et al. 2014; Olsen and Connor 2017; Levasseur et al. 2019; Zhang et al. 2019; Liang et al. 2021; Park et al. 2022). By the contrary over-expression of eIF5A in humans, mainly the EIF5A2 isoform, is observed in many cancer types, whereas correlates with metastatic processes (Jenkins et al. 2001; Caraglia et al. 2013; Wang et al. 2013; Mathews and Hershey 2015; Ning et al. 2020; Tauc et al. 2021; Nakanishi and Cleveland 2024).

*Saccharomyces cerevisiae* genome has been for long an excellent model for studying genome duplications and the driving forces and mechanisms that maintain duplicated copies of a gene. This yeast experimented an ancient whole genome duplication (WGD) event (Wolfe and Shields 1997; Kellis et al. 2004), due to an allopolyploidization (Marcet-Houben and Gabaldón 2015). However, it is estimated that only around 13% of all current yeast proteins derive from this duplication (Wolfe and Shields 1997; Seoighe and Wolfe 1998). Most of duplicated genes perform closely related functions although evolved asymmetric expression and subfunctionalization (Hittinger and Carroll 2007). In duplicates, there is an over-representation of functional categories, such as stress-response, glucose metabolism and cytosolic translation (Seoighe and Wolfe 1999; Burhans et al. 2006). It has been proposed that simple pre-WGD networks were converted into multi-stranded parallelized sub-networks by coordinated regulation of sister paralogues. Switching from one sub-network to another allows the yeast to adapt to different stresses, and to reconfigure metabolism fluxes to adapt to the available carbon source (MacKintosh and Ferrier 2018). Additionally, one of the main selective pressures to keep duplicated genes in the genome is the compensation of a loss of function in one copy by the other copy or copies. Consequently, yeast genes classified as essentials are significantly under-duplicated compared to the average for the genome (Seoighe and Wolfe 1999; Gu et al. 2003; Ihmels et al. 2007; Stein and Aloy 2008). However, in the case of eIF5A, mutations inactivating the highly expressed eIF5A gene copy (*TIF51A* in yeast) cause lethality, indicating that the second copy does not provide a classical back-up genetic compensation mechanism (Schnier et al. 1991; Schwelberger et al. 1993; Wöhl et al. 1993).

The eIF5A paralog genes form part of a specialized system in *S. cerevisiae* that responds to oxygen availability and contains several aerobic/hypoxic (anaerobic) pairs such as *TIF51A*/*TIF51B* and others encoding components of the mitochondrial electron transfer complexes, such as *CYC1/CYC7* and *COX5A/COX5B* (Zitomer and Lowry 1992). It is assumed that paralogue genes after WGD provided the basis for the capacity of *S. cerevisiae* of growing both under respiratory and fermentation-based metabolisms (Bolotin-Fukuhara et al. 2005; Fang et al. 2009). The regulation of this dual system is mostly coordinated by the oxygen-responding transcription factors Hap1, Rox1 and Mot3. Hap1 acts as an activator when bounded to a heme group, which is synthesized under oxygen availability. Heme-Hap1 activates several genes involved in the mitochondrial respiratory process including *TIF51A* and *COX5A* (Zitomer and Lowry 1992; Zhang and Hach 1999; Schüller 2003; Barba-Aliaga et al. 2020; Barba-Aliaga and Alepuz 2022b; Barba-Aliaga and Alepuz 2022a). Heme-Hap1 also binds and activates the genes encoding the transcriptional repressors Rox1 and Mot3, which in turn inhibit the transcription of hypoxic genes, such as *TIF51B* and *COX5B*. The Rox1 repressor, in synergy with Mot3, binds to the DNA through its HMG (High Mobility Group) domain, and both factors recruit the general repression complex formed by Tup1/Ssn6 to the promoter of hypoxic genes (Kastaniotis et al. 2000). By contrast, under hypoxia or anaerobiosis, heme concentration is reduced and the heme unbound form of Hap1 switches to a repressor on *TIF51A* and *ROX1* and *MOT3* promoters. Thus, reduction in Rox1 and Mot3 protein levels drives the up-regulation of *TIF51B* and other hypoxic genes (Kastaniotis et al. 2000; Sertil 2003; Lai et al. 2006; Hickman and Winston 2007; Barba-Aliaga et al. 2020; Barba-Aliaga and Alepuz 2022b).

Genetic screening of suppressors of growth defects caused by mutations in a single gene has long been used to gain insight into the function of the encoded protein. Therefore, identifying loss- or gain-of-function mutations in other genes and pathways that can compensate for the lack of the essential gene under investigation provides valuable information about the cellular processes involving our protein of interest. In the present study, we investigated the genetic mechanisms in *S. cerevisiae* that restore cell viability following the introduction of temperature-sensitive mutations in the essential gene *TIF51A*, which impair eIF5A activity. Using two genetic backgrounds, one retaining the paralogous copy (*TIF51B* background) and another deleting it (*tif51BΔ* background), we assessed how suppressor mutations arise to restore viability. Our results show that selection restores viability either by compensating for the loss of function of one paralog through the activity of the other or suppressing the temperature-sensitive degradation of Tif51A protein. These findings provide new insights into the mechanisms of genetic compensation and highlight the role of paralogs in buffering essential cellular functions within a population. Furthermore, the results suggest that no other proteins or pathways are capable of suppressing the essentiality of eIF5A.

## MATERIALS AND METHODS

### Yeast strains, plasmids, and growth conditions

All *S. cerevisiae* strains and plasmids used herein are listed in Supplementary Tables S1 and S2 respectively (see File S1). Cells were cultured in liquid YPD (2% glucose, 2% peptone, 1% yeast extract), synthetic complete media lacking uracil (SC-Ura; 2% glucose, 0.67% Yeast Nitrogen Base [YNB], 0.193% Drop-Out Mix-URA [Kaiser, Formedium]) or rich-glycerol media (YPGly; 2% glycerol, 2% peptone, 1% yeast extract). Solid media were prepared by adding 2% agar.

Experimental assays were performed with exponentially growing cells for at least four generations until the required Optical Density 600 nm (OD_600_) was reached at the corresponding temperature. For experiments involving temperature-sensitive strains, the cells were grown at 25°C (the permissive temperature) until the required OD_600_ was reached. Then, the cells were transferred to 37°C (the non-permissive temperature) for 4 or 6 h to completely deplete eIF5A (Li et al. 2014). To study the protein stability, the media were supplemented with 100 µg/ml cycloheximide (CHX) (Sigma-Aldrich), and samples were collected over 3 h. For detailed procedures of plasmids and strains generation and all materials used see Supplementary Materials and Methods in File S2.

### Isolation and characterization of spontaneous suppressor mutants

Spontaneous suppressor of the temperature-sensitive strains *tif51A-1* and *tif51A-3* in the *TIF51B* and *tif51BΔ* backgrounds were isolated by selecting for the reversion of the thermosensitive growth phenotype at 37°C. To achieve this, at least three independent cultures of each genetic background were grown at the permissive temperature of 25°C until exponential phase (OD_600_ approximately 0.5). The cells were then plated in YPD medium and the plates were incubated at 37°C for two days. Colonies exhibiting restored growth under restrictive conditions were selected as candidate suppressors. Individual isolates were restreaked onto YPD plates and grown at 37°C to confirm the stability of the phenotype. To determine the frequency of suppressor appearance, 400 µL of the undiluted culture containing approximately 10^7^ cells was plated in duplicate onto YPD plates and incubated at 37°C. Concurrently, to determine the total number of viable cells, 200 µL of a 1:10^-4^ dilution were plated onto YPD plates and incubated at 25°C. The frequency of suppressor appearance was calculated as the ratio between the c.f.u. (colony-forming units) obtained at 37°C respect to the c.f.u. obtained at 25°C.

### Genomic DNA extraction and Sanger sequencing

For genomic DNA extraction, 4 mL of an overnight culture was harvested by centrifugation and washed with sterile water. DNA was extracted from the pellets in 500 µL of Buffer 1 (50 mM Tris-HCl pH 7.5, 20 mM EDTA, 1% SDS) and using 0.5 mm glass beads and a Precellys 24 tissue homogenizer (Bertin Technologies). Genomic DNA was dissolved in TE buffer (10 mM Tris-HCl pH 7.5, 1 mM EDTA) and DNA concentration and purity were assessed with a NanoDrop spectrophotometer (Thermo Fisher Scientific). The *TIF51A* ORF was amplified by conventional PCR using the primers listed in Supplementary Table S3 (“Gene amplification for Sanger sequencing” subsection, File S1). The resulting PCR products were purified and sequenced by Sanger sequencing (Macrogen Spain).

### Whole genome sequencing (WGS) and variant calling

Genomic DNA was extracted from overnight yeast cultures grown in YPD at 25°C using a modified zymolyase–SDS protocol (Querol et al. 1992), and quality was assessed with a Qubit Flex fluorometer. WGS was performed at the FISABIO sequencing service (València) using the Illumina DNA Prep kit, with paired-end sequencing (2×150 bp) on a NextSeq 2000 platform. The sample containing indexed libraries was loaded onto NextSeq™ 2000 P1 XLEAP-SBS™ Reagent Kit (300 Cycles) (Illumina) and onto the instrument, along with the flow cell. Automated cluster generation and paired-end sequencing with dual indexes reads was performed (2×150 bp run).

Sequencing reads were quality-checked with FastQC and MultiQC, trimmed with Trim Galore, and aligned to the *S. cerevisiae* S288C reference genome (R64-1-1, (Cherry 1998)) using BWA-MEM (Li 2013). Genome-wide coverage and ploidy were assessed with an adapted version of SppIDer (Langdon et al. 2018). Variant calling followed GATK best practices (Auwera and O’Connor 2020) assuming haploid ploidy, with SNP/INDEL hard filtering and BQSR. Variants were functionally annotated with SnpEff (Cingolani et al. 2012), and suppressor-specific variants were identified based on genotype transitions using custom Python and R scripts. Further details of the protocol are provided in the Supplementary Materials and Methods (File S2).

### CRISPR/Cas9 mediated genome editing

The CRISPR-Cas9 tool was used to generate point mutations in the *ROX1* gene and an insertion in the *MOT3* gene as previously described in (Shaw and Ellis 2018). Plasmid pWS158-Cas9-*URA3* (Addgene plasmid #90517) was a gift from Dr Tom Ellis (Imperial College, UK). The sgRNA target sequences were designed using the CHOPCHOP web tool (https://chopchop.cbu.uib.no) and assembled to the Cas9-sgRNA expression vector pWS158. Repair templates (donor DNA) providing the desired mutations were synthesized by PCR. All the primers used for sgRNA and donor DNA are listed in Table S3 (“Genome editing by CRISPR-Cas9” subsection). This strategy was used to introduce point mutations in *ROX1* [G158T (SNP1), G192T (SNP3), and T218C (SNP2)] and a 17-pb insertion (nucleotides 697–714) in *MOT3*, that causes a frameshift mutation. The corresponding strains were transformed following the lithium acetate-based method (Shaw and Ellis 2018) and transformants were selected in SC-Ura. The loss of thermosensitivity was subsequently confirmed, and the introduced mutations were confirmed by Sanger sequencing (Macrogen Spain). For detailed procedures see Supplementary Materials and Methods (File S2).

### Phenotypic assays

For spot assays, cells were grown in YPD medium at 25°C and harvested during the mid-exponential growth phase. 5-fold serial dilutions were prepared, starting at an OD_600_ of 0.3 and 5 µL aliquots were spotted onto YPD, YPGly, or SC-Ura plates. Plates were incubated at 25°C, 30°C, and 37°C for 2 – 3 days.

For growth curve analysis in microtiter plates, cells were pre-cultured in deep 96-well plates with 500 µL of YPD, YPGly or SC-Ura media at 25°C until saturation. After pre-culture, the OD_600_ was measured and adjusted to 1.0, and 20 µl of cell suspension were inoculated into 96-well plates (Thermo Fisher Scientific) to a final volume of 200 µL per well in YPD, YPGly or SC-Ura media (initial OD_600_ of 0.1). To monitor growth under different conditions, the plates were incubated in a SPECTROstar Nano (BMG LABTECH) plate reader at 25°C, 30°C or 37°C. Absorbance at 600 nm was recorded every 30 min for 24 – 48 h under static conditions and the experiments were concluded once the strains reached stationary phase. Growth rates were estimated from the exponential growth phase by linear regression of the natural logarithm of OD_600_ values (ln OD_600_) versus time. The specific growth rate was calculated as the slope of the regression line. At least three biological replicates were analyzed for each strain. For detailed procedures see Supplementary Materials and Methods (File S2).

### Western blotting

For protein analysis by western blotting, we followed the protocol described by (Zuzuarregui et al. 2015). Samples were run on SDS-PAGE gels, transferred, and immunoprobed as described in the Supplementary Materials and Methods (File S2). The antibodies used in this study are listed in Supplementary Table S4 (File S1). To capture variation across all samples, the signal in each lane was normalized to the mean signal across all lanes in a single blot. Then, the resulting signal of the bands was normalized against the corresponding G6pdh signal. At least three biological replicates of each sample were analyzed for quantification purposes; however, cycloheximide-based assays were performed as a single replicate for qualitative assessment. Further details of the protocol are provided in the Supplementary Materials and Methods (File S2).

### RT-qPCR analysis

To analyze the mRNA levels, total RNA was isolated from yeast cells using the protocol described in (Barba-Aliaga and Alepuz 2022b). Reverse transcription and quantitative PCR reactions were performed as detailed in (Garre et al. 2013). Relative mRNA levels were normalized to the endogenous *ACT1* control. At least three biological replicates of each sample were analyzed. The specific primers used for gene amplification are listed in Supplementary Table S3 (File S1). More details can be found in the Supplementary Materials and Methods (File S2).

### Chromatin immunoprecipitation (ChIP)

The chromatin immunoprecipitation (ChIP) experiments were performed as previously described (Li et al. 2016) with minor modifications. qPCR was run as described above using the primers listed in Supplementary Table S3 (“Gene expression detection by RT-qPCR and ChIP-qPCR” subsection). The qPCR amplification data were normalized with the total INPUT DNA value in the corresponding whole cell extract. Immunoprecipitations of RNA Polymerase II (RNA pol II) were made with anti-Rpb1 antibody (8WG16; Invitrogen (MA1-10882)) and Dynabeads Pan Mouse IgG (Invitrogen (11041)); and for Rox1-13myc with anti-myc antibody (Invitrogen (13-2500)) and Dynabeads Pan Mouse IgG (Invitrogen (11041)). At least three biological replicates of each sample were analyzed. More detailed procedures can be found in the Supplementary Materials and Methods (File S2).

### Protein sequence alignments, structural modeling and protein stability prediction

Amino acid sequences for *S. cerevisiae* Tif51A (eIF5A1), Tif51B (eIF5A2) and its respective orthologs were retrieved from the Saccharomyces Genome Database (SGD) and UniProt. Multiple sequence alignments were performed using the Clustal Omega algorithm via the UniProt alignment service. The resulting alignments were imported into Jalview v2.11.5.1 (Waterhouse et al. 2009) for visualization, conservation analysis, and figure generation. To evaluate protein conservation, pairwise sequence alignments were performed with EMBOSS Needle (PSA) (https://www.ebi.ac.uk/jdispatcher/psa/emboss_needle; (Madeira et al. 2024)), and the percentage of sequence identity of Tif51A (eIF5A1) and its respective orthologs was calculated.

To assess the structural impact of the identified mutations, three-dimensional (3D) models of wild-type and mutant variants of Tif51A and Rox1 were predicted using AlphaFold2 via the ColabFold pipeline v1.5.5 (Mirdita et al. 2022). For each protein, the best-ranked model based on the predicted local distance difference test (pLDDT) score was selected for further analysis. Molecular graphics and analyses to compare conformational changes between the wild-type and mutated variants were performed with UCSF Chimera v1.18 (Pettersen et al. 2004).

The effect of single-site mutations on protein stability was predicted using the I-Mutant2.0 server (https://folding.biofold.org/i-mutant/i-mutant2.0.html; (Capriotti et al. 2005)), based on the protein sequence. Stability changes were evaluated by calculating the free energy change ΔΔG at 25°C and pH 7.0.

### Statistical analysis

Statistical analyses were performed using GraphPad Prism 8 (GraphPad Software). Differences between two groups were analyzed using a two-tailed unpaired Student’s t-test, whereas comparisons among multiple groups were performed using one-way analysis of variance (one-way ANOVA) followed by Bonferroni’s multiple comparisons test. A minimum of three biological replicates was analyzed for each experiment.

## RESULTS

### Suppressors of temperature-sensitive *TIF51A* mutations in *S. cerevisiae* show up-regulation of the *TIF51B* gene

To gain further knowledge about the molecular functions of the evolutionary conserved eIF5A protein, we conducted a genetic screening to isolate suppressors that counteract the growth impairment caused by mutations in the essential eIF5A-encoding gene *TIF51A* in *S. cerevisiae*. For this we used the temperature-sensitive strains *tif51A-1* and *tif51A-3*, which contain one (Pro83 to Ser) and two (Cys39 to Tyr and Gly118 to Asp) point mutations in eIF5A protein, respectively (Li et al. 2014). We grew the two strains, *tif51A-1* and *tif51A-3*, in glucose-rich YPD medium at 25°C until exponential phase. We then plated onto YPD plates and incubated them at 25°C and 37°C for two days (see Materials and Methods section for details). We observed the growth of a few suppressor colonies from each strain at 37°C. The determination of the frequency of spontaneous suppressor emergence showed similar values for both strains, which were consistent with loss-of-function mutations (frequencies of around 10^-6^). We selected four suppressor clones of each strain for further characterization (Fig. 1a).

**Figure 1.**
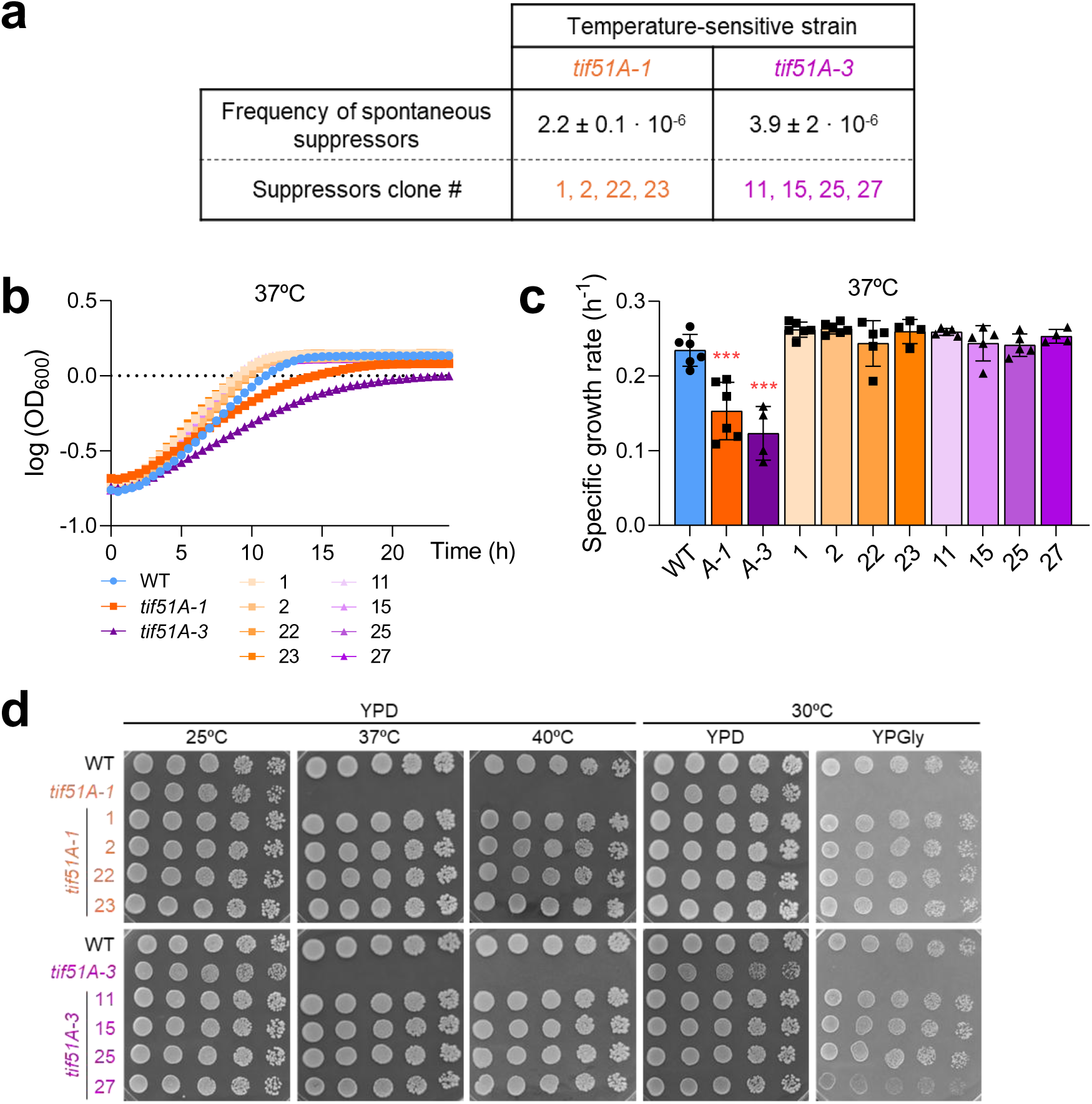
Isolation of suppressors of temperature-sensitive mutations in the yeast eIF5A encoding gene *TIF51A*. **(a)** Frequency of spontaneous suppressor emergence. Frequencies were determined by growing *tif51A-1* and *tif51A-3* yeast cells in YPD to mid-exponential phase at 25°C and plating onto YPD agar. Plates were incubated at 25°C (permissive) and 37°C (restrictive) for 2 days. The frequency of suppressor appearance was calculated as the ratio between the c.f.u. (colony-forming units) obtained at 37°C respect to the c.f.u. obtained at 25°C. Specific independent suppressor clones selected for further characterization are identified by the numbers in orange (*tif51A-1* background) and purple (*tif51A-3* background). **(b, c)** Growth kinetics of suppressor strains at restrictive temperature. Wild-type (WT), *tif51A-1* and *tif51A-3* strains, along with their respective suppressors, were cultured in YPD at 25°C. Cultures were then diluted to an OD_600_ of 0.1 and shifted to 37°C. Growth was monitored by recording OD_600_ every 30 min for 24 h using a SPECTROstar Nano plate reader. Growth curves (b) are represented as the mean of log_10_ (OD_600_) from at least three biological replicates, and specific growth rates (c) were calculated from the slope of the exponential phase. Statistical significance was determined using a one-way ANOVA relative to the wild-type strain followed by Bonferroni’s post hoc multiple-comparison test. ***p < 0.001. Growth curves with error bars are provided in supplementary Figure S1. **(d)** Spot assay of suppressor strains. Same yeast strains used in (b, c) were cultured in YPD at 25°C and 5-fold serial dilutions were spotted onto YPD and YPGly plates. Plates were incubated for 2 – 3 days at 25°C, 37°C, and 40°C to assess temperature sensitivity, and at 30°C on YPGly to evaluate respiratory capacity.

Next, we studied the growth of the suppressors at different conditions. First, we monitored cell growth at permissive (25°C) and restrictive temperatures (37°C). All suppressors grew similar to the wild-type strain in YPD at 37°C (Fig. 1b and c and Supplementary Fig. S1). To study the extension of the suppression of the *TIF51A* essentiality, we grew the suppressors under more restrictive conditions. For this we incubated cells in glucose rich media at 40°C and in media with glycerol as the only carbon source at 30°C. In this last condition, eIF5A has been proved to be essential as it is necessary for mitochondrial respiration (Barba-Aliaga et al. 2020; Barba-Aliaga and Alepuz 2022b; Barba-Aliaga et al. 2024). As shown in Fig. 1d, all suppressors grew well at 40°C and all, except sup. 27, were able to restored growth in glycerol of the strains with mutations in *TIF51A*.

We further analyzed the suppressors of the Tif51A temperature-sensitive mutations. For this, we studied the levels of Tif51A protein in parental and suppressor strains in Western-blots with protein samples obtained under permissive (25°C) and restrictive (37°C) temperature. Using an anti-Tif51A antibody, which does not recognize Tif51B protein, we observed that Tif51A protein levels were very low after incubation at 37°C in all suppressor analyzed, similar to the parental Tif51A-1 and Tif51A-3 mutant proteins (Fig. 2a). However, using an anti-hypusinated-eIF5A antibody which does recognize either hypusinated Tif51A or Tif51B protein, we observed for all suppressors a band of the expected size compatible with the Tif51B protein, as it was coincident with the Tif51A-1 band, but distinguishable of the Tif51A-3 band, which migrates with lower mobility as previously described (Li et al. 2014) (Fig. 2a). To confirm that the Tif51A-1 and Tif51A-3 proteins remained unstable at 37°C in the suppressors, while a stable Tif51B protein was detected, we conducted experiments involving translational shut-off with cycloheximide (CHX) in some of the suppressors to measured protein stability. As seen in Fig. 2b, Tif51A-1/-3 proteins were unstable and the corresponding signal was not detected after 1.5-2.0 hours of incubation at 37°C. However, a stable Tif51B protein, distinguishable in the *tif51A-3* background strains, was detected using anti-hypusinated-eIF5A antibody (Fig. 2b).

**Figure 2.**
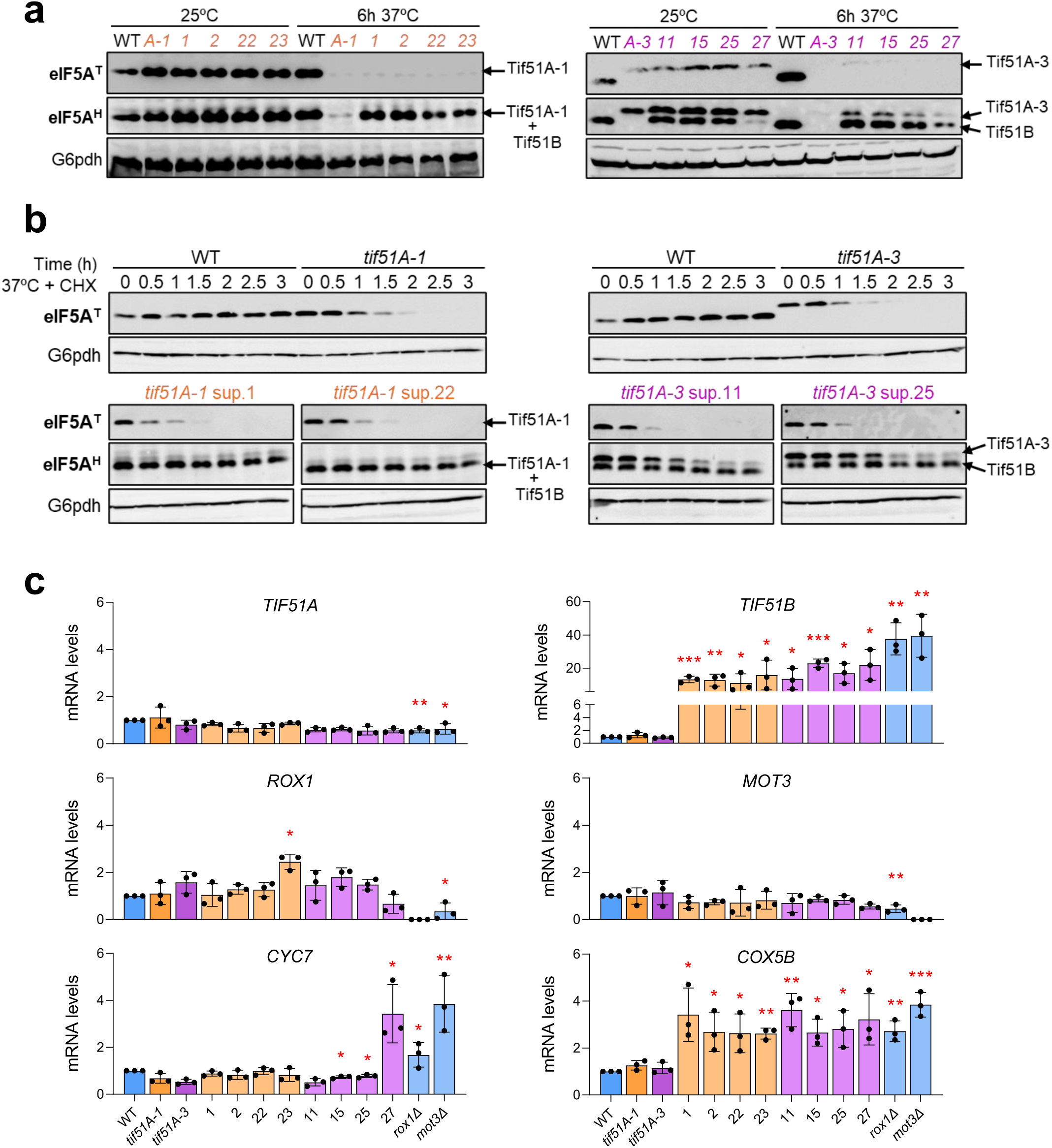
Suppressors restore the function of eIF5A by upregulating the *TIF51B* paralog gene. **(a)** Wild-type (WT), *tif51A-1*, *tif51A-3*, and the indicated suppressors were cultured in YPD at 25°C to mid-exponential phase at 25°C, and then shifted to 37°C for 6 h. Tif51A and Tif51B protein levels were determined by Western blotting using an anti-eIF5A antibody (which detects total Tif51A; eIF5A^T^), and an anti-hypusine antibody (which detects hypusinated Tif51A and Tif51B; eIF5A^H^). G6pdh protein levels were used as a loading control. **(b)** Same yeast strains used in (a) were cultured in YPD to mid-exponential phase at 25°C, then shifted to 37°C for 3 h in the presence of 100 µg/mL CHX to inhibit *de novo* protein synthesis. Tif51A and Tif51B protein levels were determined by Western blotting using same antibodies and loading control as in (a). A representative image is shown. **(c)** Wild-type (WT), the *tif51A-1*, *tif51A-3* mutants and their respective suppressors, and *rox1Δ* and *mot3Δ* mutants, were cultured in YPD to mid-exponential phase at 25°C. mRNA levels of the indicated genes were determined by RT-qPCR using ORF-specific primers, and normalized to *ACT1* mRNA levels. Data represent the mean ± SD of three biological replicates. Statistical significance was determined using a two-tailed unpaired Student’s t-test relative to the corresponding parental strain. *p<0.05, **p<0.01, ***p<0.001.

The above results were compatible with mutations that eliminate the repression of the *TIF51B* gene under aerobic conditions, potentially through alterations in its promoter or through mutations affecting the *TIF51B* repressors Rox1 and Mot3 themselves. To test the first option, we sequenced the promoter regions of *TIF51B* in several suppressors, but all sequences were identical to the native one. Then, to check whether Rox1/Mot3 had lost their repressor capacity, we investigated the mRNA levels of *TIF51B* gene and of other duplicated genes which have a paralog described to be expressed under hypoxic conditions and repressed under aerobic conditions by Rox1/Mot3 (Lowry and Zitomer 1988; Trueblood and Poyton 1988; Sertil 2003). For this, we choose the hypoxic *CYC7* and *COX5B* genes, encoding subunits of the mitochondrial cytochrome c (Lowry and Zitomer 1988; Trueblood and Poyton 1988; Kastaniotis et al. 2000). Accordingly to the detection of Tif51B protein by Western-blot (Fig. 2a and b), a higher level of *TIF51B* mRNA was detected in all suppressor but not in the parental strains (Fig. 2c), and the levels were similar to those observed in the *rox1Δ* and *mot3Δ* deletion strains, confirming the derepression of the gene. However, the results with the other two hypoxic genes were not conclusive since *COX5B* gene seems to be derepressed in suppressors and *rox1Δ* and *mot3Δ* mutants, but *CYC7* was only observed in the suppressor 27 and in *rox1Δ* and *mot3Δ* (Fig. 2c). Therefore, we could not assume that Rox1 and Mot3 had lost their repression capacity.

Altogether, we concluded that it was possible to obtain suppressors of the temperature-lethal phenotype associated with mutations in the *TIF51A* gene, which encodes eIF5A, and that most of these suppressors restored growth at high temperatures and under respiratory conditions. Furthermore, we concluded that all the isolated suppressors show derepression of the second copy gene of eIF5A, *TIF51B*. However, this was not due to mutations in the promoter/5’UTR of *TIF51B* gene/mRNA and not to a global loss of Rox1/Mot3 function. Interestingly, these results suggest the existence of a previously undescribed unconventional backup mechanism for *TIF51A* mutations based on the paralogue *TIF51B*, which is silent under normal conditions.

### Suppressors of the temperature-sensitive *TIF51A* mutations contain mutations in Rox1 and Mot3 that derepress *TIF51B*

Because we did not observe a global derepression of genes under the control of Rox1 and Mot3 repressors under aerobic conditions, we hypothesized that suppressors could contain a mutation in another unknown protein which will specifically disturb the regulation of *TIF51B* gene. To identify that mutation/s we whole-genome sequenced all suppressor strains (see Materials and Methods section for full description). For most suppressors we found few single nucleotide polymorphisms (SNP) or deletions/insertions (INDELs) in different genes (Fig. 3a). However, all suppressors contained mutations in *ROX1* or *MOT3*: SNPs in *ROX1* for the sups. 1, 2, 23 and 25; INDELs in *ROX1* for sups. 22, 11 and 15; and INDEL in *MOT3* for sup. 27 (Fig. 3a). Therefore, we hypothesized that these mutations were yielding a selective loss of Rox1 and Mot3 repressive activity at the *TIF51B* promoter (Fig. 3b).

**Figure 3.**
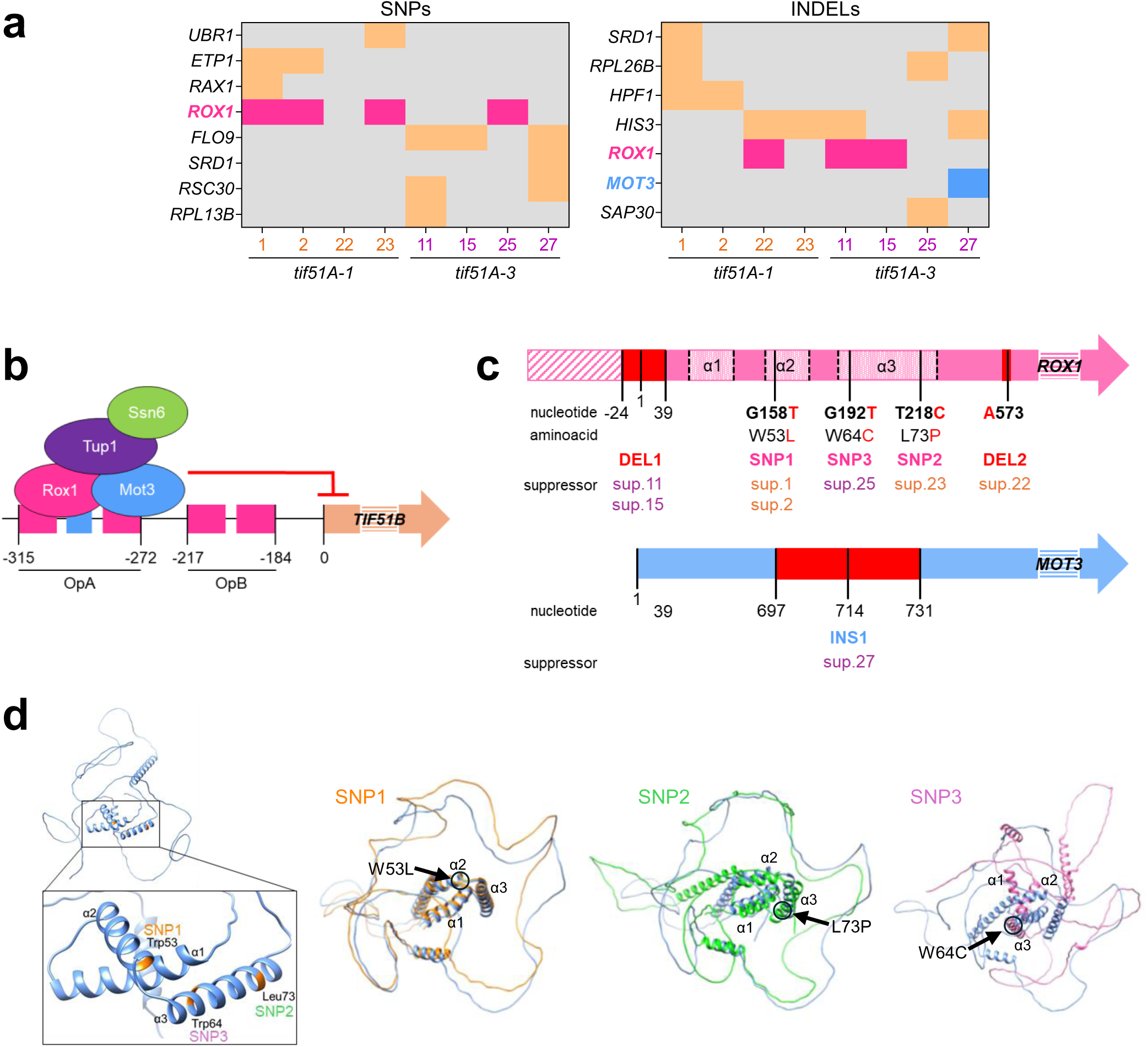
Suppressors of *TIF51A* mutations contain mutations in the genes encoding Rox1/Mot3 repressors. **(a)** Heatmaps summarizing the identified mutations (single nucleotide polymorphisms, SNPs; deletions or insertions, INDELs) across the isolated suppressors (1, 2, 22, 23, 11, 15, 25, 27) of *tif51A-1/-3* mutations in strains. **(b)** Schematic representation of the Rox1-Mot3-Tup1-Ssn6 corepressor complex binding to the *TIF51B* promoter under aerobic conditions. Boxes in the promoter indicate the Rox1 binding sites (in fuchsia) and the Mot3 binding site (in blue), which are included in the operators A and B. Numbers indicate the nucleotide position relative to the translation start site (ATG, +1). **(c)** Schematic representation of the mutations identified in *ROX1* and *MOT3*, showing the locations of deletions (DEL), SNPs, and insertions (INS) found in specific suppressor strains. **(d)** AlphaFold2-predicted models of Rox1 and the HMG-box domain. (Left) WT Rox1 with a magnified view of the HMG-box showing the amino acid residues (Trp53, Trp64, Leu73) mutated in suppressors 1, 2, 22 and 23. (Right) Superposition of models for WT Rox1 (blue) and the SNP1 (orange), SNP2 (green), and SNP3 (magenta) mutants.

To gain insight into this, we determined the structural changes in the repressor proteins that may cause the mutations identified in the suppressors. For this, we predicted the 3D-protein structure using AlphaFold2 via the ColabFold pipeline, and then, we visualized and compared the structures with UCSF Chimera (see Materials and Methods for more information). Rox1 repressor belongs to the HMG family of transcription factors and contains three DNA-binding alpha-helix in the N-terminal part of the protein (Deckert et al. 1995) (Fig. 3c and d). Sups. 1 and 2 contained the amino acid changes Trp53 to Leu (SNP1) in the α2-helix that did not modify the overlapping of the α2-helix 3D-structures between the mutant and the native Rox1 protein, but it did so in the surrounding protein regions (Fig. 3d). Sup. 25 contained the amino acid change Trp64 to Cys (SNP3) and sup. 23 contained the change Leu73 to Pro (SNP2), both in the α3-helix of Rox1 (Fig. 3c and d). The predicted protein structures of SNP2 and SNP3 showed the loss of overlapping between the 3D-protein structures of these point mutants and the native Rox1 protein both in the α-helixes and rest of the protein (Fig. 3d). The rest of suppressors contained insertions or deletions in *ROX1* or *MOT3* sequences: deletion of the first amino acids of Rox1 in sups. 11 and 15 (DEL1 with deletion from -24 to +39 nucleotides); a nucleotide deletion that altered the translation frame and inserts a stop codon in frame in sup. 22 (DEL2 with deletion of nucleotide 573); and sup. 27 that was the only one with a mutation in gene encoding the repressor Mot3, that consisted in a duplication of 17 nucleotides that also altered the translation frame and introduced soon after a stop codon (INS1) (Fig. 3c). From these data, we hypothesized that SNP1 (sup. 1 and 2), SNP2 (sup. 23) and SNP3 (sup. 25) would alter the binding of Rox1 to *TIF51B* promoter because produce important changes in the Rox1 DNA-binding motif or close regions. However, DEL2 (sup. 22) and INS1 (sup. 27) will result in truncated Rox1 and Mot3 proteins, respectively, unable to act as repressors over *TIF51B*. Meanwhile, mutation DEL1 (sup. 11 and 15) will result in no translation of the *ROX1* gene.

In order to investigate this, some of the identified mutations were introduced into the parental strains employed in the suppressor screening, using the CRISPR technique developed for yeast (Shaw and Ellis 2018). For Rox1 mutations, we used parental strains containing a C-terminal *ROX1-13myc* tag, so these strains could be used to study further the effect of the mutations in Rox1 transcriptional repression activity. Thus, *ROX1* mutations SNP1 and SNP2 were introduced into the *tif51A-1* strain and SNP3 mutation into the *tif51A-3*, both containing *ROX1-13myc.* Additionally, we introduced INS1 mutation of *MOT3* gene in the parental *tif51A-3.* First, we observed that the three *ROX1* SNP mutants and *MOT3* INS1 grew well at permissive temperature (25°C) and suppressed the grow defect of the parental *tif51A* mutants at restrictive temperature (37°C) (Fig. 4a). Then, we determined the *TIF51B* mRNA levels in all strains and observed higher levels in *ROX1* SNP1, SNP2, SNP3 and *MOT3* INS1 mutants than in the corresponding *tif51A-1/-3* parental and the wild-type strains (Fig. 4b). Confirming that higher *TIF51B* mRNA levels are due to the derepression of the gene, we observed a higher binding of the RNA polymerase II to *TIF51B* in the *ROX1* SNPs and *MOT3* INS1 mutants by chromatin immunoprecipitation assays (ChIP) (Fig. 4c). We next investigated whether *TIF51B* derepression was due to a reduced expression of the repressors when containing the mutations or whether the mutations impair the binding of the repressors to *TIF51B* promoter. We observed that Rox1 proteins containing the amino acid changes due to the SNPs mutations were expressed at similar levels than native Rox1 protein (Fig. 4d). However, ChIP assays showed that Rox1 binding to the *TIF51B* promoter was lost in the SNP1, SNP2 and SNP3 mutant versions (Fig. 4e).

**Figure 4.**
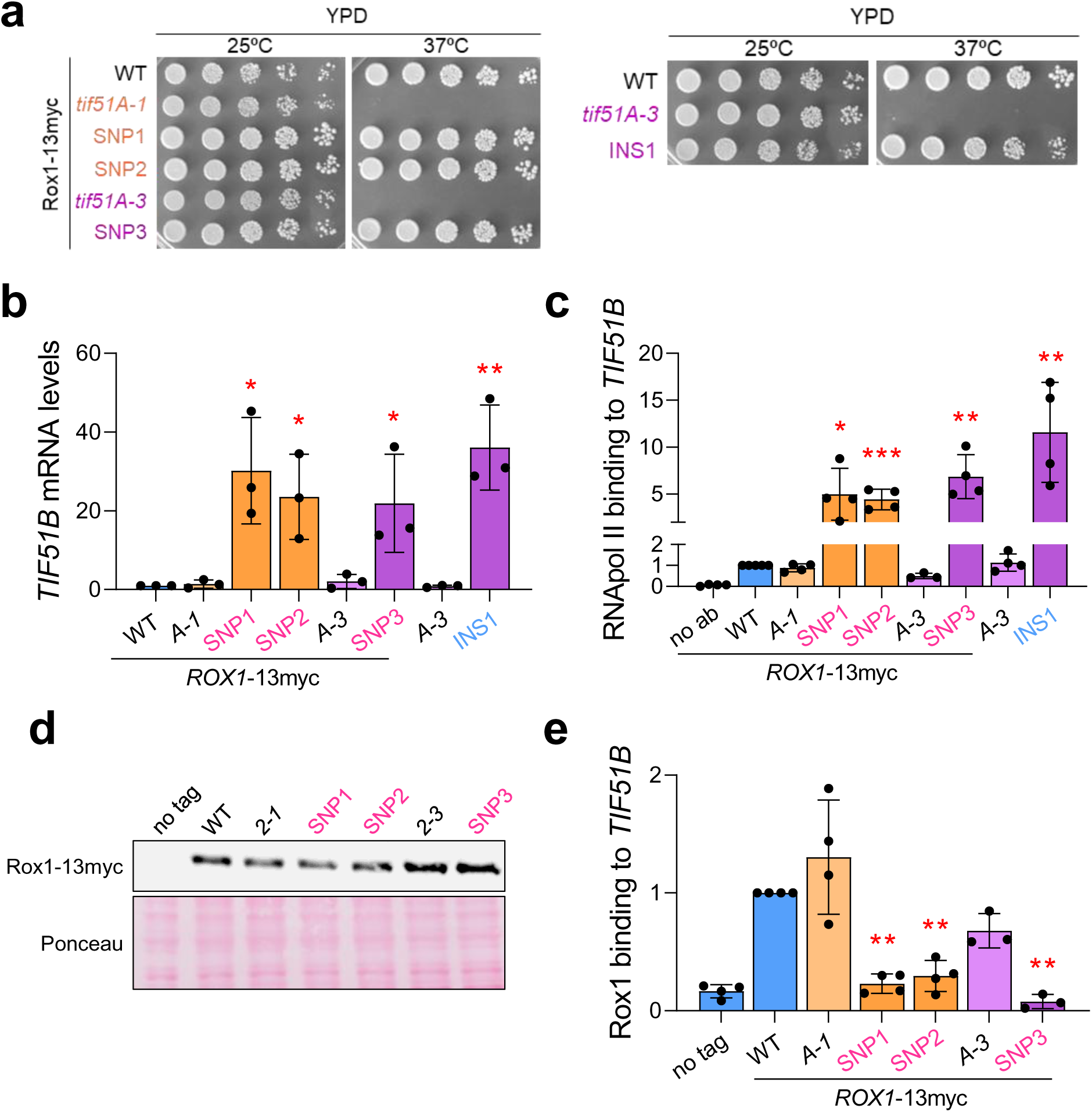
Mutations in the Rox1 and Mot3 repressors prevent their binding to the *TIF51B* promoter and, therefore, impair their repressive function. **(a – e)** Strains carrying a genomic *ROX1* C-terminal myc tag (*ROX1*-13myc), including wild-type (WT), *tif51A-1*, *tif51A-3* and SNP1-3 strains, as well as the non-tagged *tif51A-3* and INS1 strains, were cultured in YPD to mid-exponential phase at 25°C. Then, cells were collected for subsequent analysis. **(a)** A spot assay was performed by spotting 5-fold serial dilutions of the cell cultures onto YPD plates. Plates were incubated for 2-3 days at 25°C and 37°C to assess temperature sensitivity. **(b)** mRNA levels of *TIF51B* were determined by RT-qPCR using ORF-specific primers, and normalized to *ACT1* mRNA levels. **(c, e)** Binding of RNA Polymerase II (RNApol II) to *TIF51B* promoter was determined by ChIP assays using an anti-Rpb1 antibody (8WG16) **(c)** and of Rox1-13myc using an anti-myc antibody **(e)**. Immunoprecipitated DNA was used to quantify the binding using *TIF51B* ORF-specific primers **(c)** and promoter-specific primers **(e)**. Results are expressed as the percentage of the signal obtained in each ChIP sample relative to the signal obtained with the DNA from the corresponding whole-cell extract (input). A sample without antibody (no ab) was used as a negative control. **(d)** Rox1-myc protein levels were determined by Western blotting using an anti-myc antibody. Ponceau staining is shown as a loading control. **(b, c, e)** Data represent the mean ± SD of at least three biological replicates. Statistical significance was determined using a two-tailed unpaired Student’s t-test relative to the corresponding parental strain. *p < 0.05, **p < 0.01, ***p < 0.001.

Overall, our results show that all the isolated suppressors of the temperature-sensitive *TIF51A* mutations contained mutations (SNPs or INDELs) in the genes that encode Rox1 or Mot3 repressors. Furthermore, our results demonstrate that introducing any of the three SNP mutations in the α-helix 2 and 3 of Rox1 impairs its binding to the *TIF51B* promoter, resulting in the expression of the Tif51B protein, which compensates for the absence of the Tif51A-1 and Tif51A-3 temperature-sensitive proteins at high temperatures.

### Repression by *ROX1* and *MOT3* acts differentially over anaerobic paralogs according to the distribution of their binding sites within the gene promoters, but maintains a strict *TIF51B* repression to prevent the deleterious effect of eIF5A over-expression

To gain a deeper understanding on the regulation of other gene pairs controlled in opposite ways by aerobic/anoxic conditions through the Rox1/Mot3 repressor system (Lowry and Zitomer 1988; Trueblood and Poyton 1988; Hodge et al. 1989; Sertil 2003), such as *CYC1/CYC7* and *COX5A/COX5B* pairs, and because we observed different regulation patterns of the anoxic copies *CYC7* and *COX5B* in the isolated suppressors (see Fig. 2c), we studied their regulation in the strains containing *ROX1* SNP1/2/3 and *MOT3* INS1 mutants. As seen in Fig. 5a, *COX5B* gene was derepressed at similar levels in strains with SNP1, SNP2 and SNP3 mutations in *ROX1* and with INS1 mutation in *MOT3*; whereas *CYC7* expression was not affected in the *ROX1* SNP mutants but was derepressed in the *MOT3* INS1 mutant, accordingly with previous results obtained with the isolated suppressors (Fig. 5a and Fig. 2c). To understand the results, we searched for all putative binding sites of Rox1 and Mot3 in the promoter sequences using the previously proposed consensus binding sequences YYHATTGTTCTC for Rox1 (Lowry et al. 1990) and TRCCTD for Mot3 (Grishin et al. 1998) where Y indicates a pyrimidine nucleotide (nt), H indicates that G nt is the only base missing, R indicates a purine nt and D indicates that C nt is the only base missing (Fig. 5b and Supplementary Table S5). *TIF51B* promoter contains 4 putative Rox1 and 2 putative Mot3 binding sites accordingly to the strong repression made through the Rox1 and Mot3 repressors. However, the anoxic gene *CYC7* contains 2 putative degenerated Rox1 and 2 putative Mot3 binding sites in its promoter; and *COX5B* promoter contains 2 and 1 canonical sequences for the binding of Rox1 and Mot3, respectively. The differential distribution of the repressor sites in these two genes and the lack of the Rox1 core consensus binding sequence in the case of *CYC7* explains, therefore, the regulation observed that in the case of *CYC7* only depends on Mot3 but in *COX5B* gene depends to a similar extend on Rox1 and Mot3 (Fig. 5a and b).

**Figure 5.**
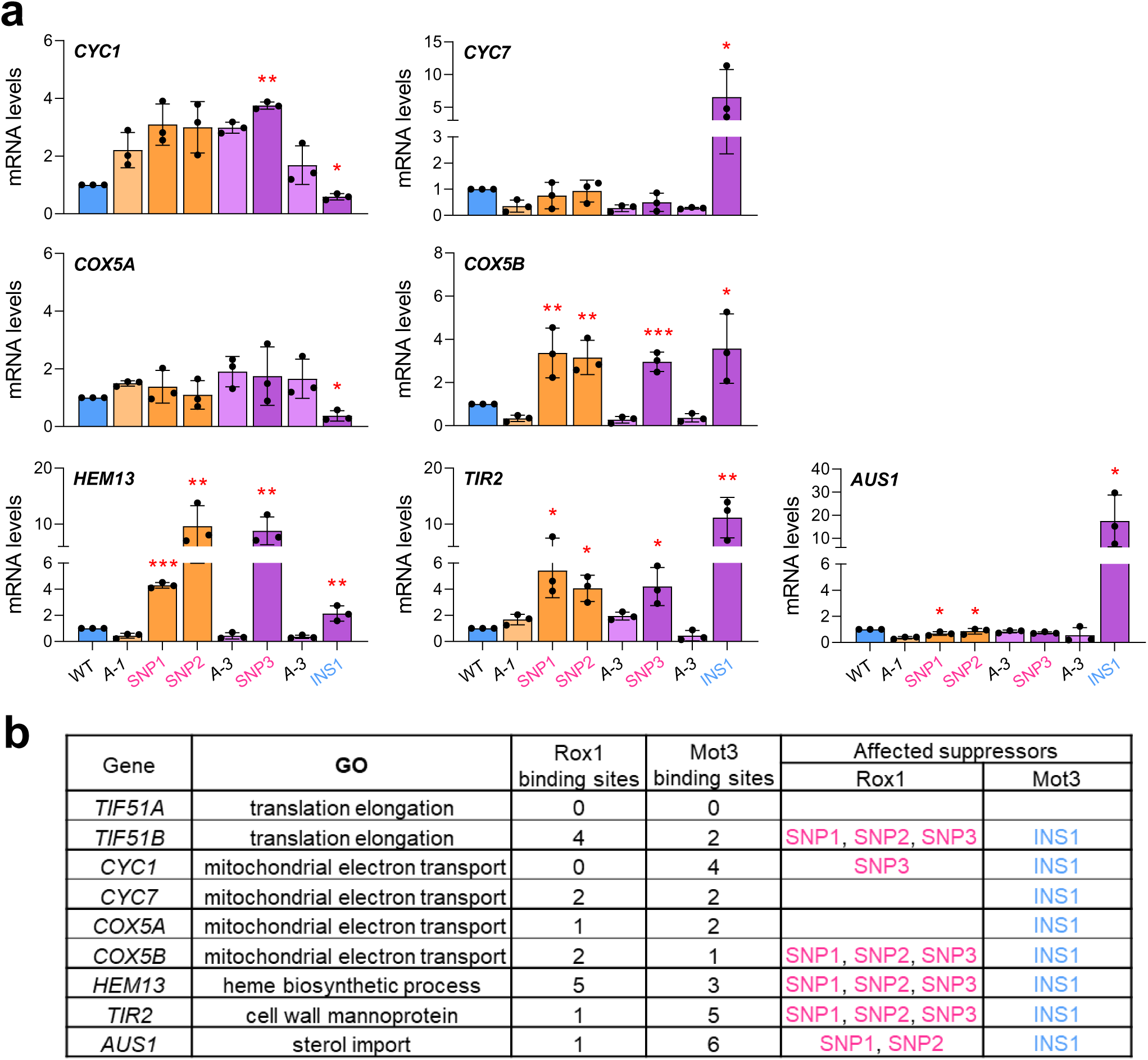
Regulation of normoxic/hypoxic genes by the Rox1 and Mot3 repressors correlates with the relative number of their respective binding sites at gene promoters. **(a)** Strains carrying a genomic *ROX1* C-terminal myc tag (*ROX1*-13myc), including wild-type (WT), *tif51A-1*, *tif51A-3* and SNP1-3 strains, as well as the non-tagged *tif51A-3* and INS1 strains, were cultured in YPD to mid-exponential phase at 25°C. mRNA levels of the indicated genes were determined by RT-qPCR using ORF-specific primers, and normalized to *ACT1* mRNA levels. Data represent the mean ± SD of at least three biological replicates. Statistical significance was determined using a two-tailed unpaired Student’s t-test relative to the corresponding parental strain. *p < 0.05, **p < 0.01, ***p < 0.001. **(b)** Summary table of normoxic/hypoxic genes is shown, indicating Gene Ontology (GO) terms, the number of putative Rox1 and Mot3 binding sites in the respective promoters, and the specific suppressors where these genes are significantly up- or down-regulated. The sequence and position of the putative Rox1 and Mot3 biding sites in the gene promoters can be seen in Supplementary Table S5.

To corroborate the correlation in Rox1/Mot3 repression and their relative number of binding sites in promoters, we additionally studied other genes repressed by these factors. We choose *HEM13*, encoding a protein involved in the biosynthesis of heme groups (Urban-Grimal and Labbe-Bois 1981; Lowry et al. 1990; Klinkenberg et al. 2005), and whose promoter contains a high number of Rox1 (5) and Mot3 (3) putative binding sites; *TIR2*, encoding a cell wall mannoprotein (Kowalski et al. 1995; Abramova et al. 2001; Kwast et al. 2002) and whose promoter contains more Mot3 binding sites (5) than Rox1 binding sites (1); and *AUS1*, encoding a protein of the sterol import system (Wilcox et al. 2002; Alimardani et al. 2004; Sullivan et al. 2009) and with also more putative binding sites for Mot3 (6) than for Rox1 (1) in its promoter. Accordingly, *HEM13* was derepressed in strains with *ROX1* SNP mutations and to a lower extend in *MOT3* INS1 mutant (Fig. 5b and Supplementary Table S5). By the contrary, *TIR2* and *AUS1* genes showed high derepression in *MOT3* INS1 mutant and lower or no derepression, respectively, in *ROX1* SNP mutants (Fig. 5a and b and Supplementary Table S5). In sum, the results obtained demonstrate that there is a specific and differential regulation of the anoxic genes controlled by Rox1 and Mot3 that depends on the relative number of their binding sites in the gene promoters. According to the results, the *TIF51B* gene is one with higher number of Rox1 and Mot3 binding sites and stronger induction in the cells with the described mutations in the two repressors (Fig. 4b and 5a).

The results above, in conjunction with previous works, demonstrate that the second copy gene encoding eIF5A, *TIF51B*, is under strict repression control provided by the two repressors Rox1 and Mot3. This led to the hypotheses that exogenous expression of *TIF51B* could be deleterious for cells under aerobic conditions when *TIF51A* is highly expressed. This hypothesis was tested by the introduction of plasmids over-expressing Tif51A, Tif51B, or a control empty plasmid, in the wild-type strain. The effect of these plasmids on growth at 25°C and 37°C was then examined. Wild-type cells exhibited a slight decline in growth at both temperatures when additional doses of eIF5A were added by expressing *TIF51A* or *TIF51B* under the control of the strong constitutive *TEF* promoter, in comparison to control cells with an empty plasmid (Fig. 6). Thus, the results suggest that a strong repression of the *TIF51B* gene impairs the detrimental effect of eIF5A over-expression.

**Figure 6.**
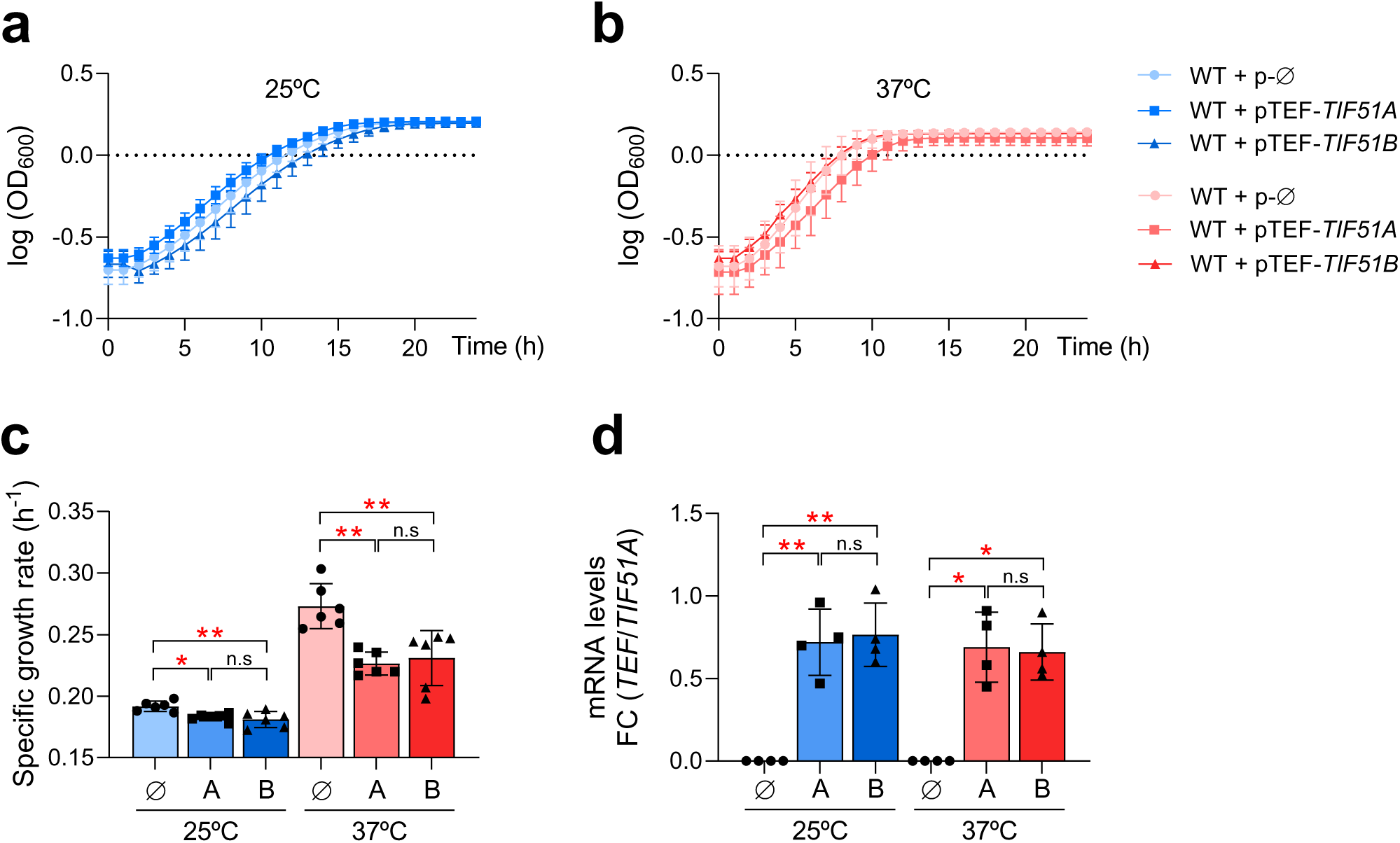
Over-expression of eIF5A has a detrimental effect on yeast cell growth. **(a – c)** Wild-type (WT) strain carrying either the empty vector pRS416-URA3 (p-Ø) or plasmids pRS416-*TEF-TIF51A* (pTEF-*TIF51A*) or pRS416-*TEF-TIF51B* (pTEF-*TIF51B*), were cultured in SC-Ura to mid-exponential phase at 25°C. Cultures were then diluted to an OD_600_ of 0.1 and either maintained at 25°C (a) or shifted to 37°C (b). Growth was monitored by recording the OD_600_ every hour for 24 h using a SPECTROstar Nano plate reader. Growth curves (a, b) are represented as the mean ± SD of log_10_ (OD_600_) from at least four biological replicates, and specific growth rates (c) were calculated from the slope of the exponential phase. **(d)** mRNA expression levels of the strains described in (a – c). Strains were cultured in SC-Ura to mid-exponential phase at 25°C, and then shifted to 37°C for 4 h. mRNA levels of endogenous *TIF51A* and plasmid-derived *TEF-TIF51A/B* were determined by RT-qPCR using specific primers, and normalized to *ACT1* mRNA levels. Data are expressed as the ratio of plasmid-derived mRNA relative to endogenous *TIF51A* mRNA levels (*TEF/TIF51A* fold change). Data represent the mean ± SD of four biological replicates. **(c, d)** Statistical significance was determined using a one-way ANOVA followed by Bonferroni’s post hoc multiple-comparison test. *p < 0.05, **p < 0.01; n.s, non-significant.

### Suppressors of the temperature-sensitive *TIF51A* mutations in yeast cells with *tif51BΔ* background are revertants or new mutations in Tif51A protein that confer temperature stability

Because all the suppressors isolated in our screening allowed the growth of temperature-sensitive *TIF51A* mutants at the restrictive temperature through the upregulation of the second eIF5A gene copy, *TIF51B*, we decided to conduct another screening of suppressors of the *TIF51A* mutants in yeast cells in which *TIF51B* was deleted. The aim of this experiment was again to identify new mutations in genes and/or pathways that compensate for the absence of the eIF5A protein in cells, but avoiding suppression through the up-regulation of the second copy of the eIF5A gene. To do so, we followed the same protocol for the isolation of suppressors described above, but working with the yeast strains *tif51A-1 tif51BΔ* and *tif51A-3 tif51BΔ*. We observed the growth of few suppressor colonies of each strain at 37°C. The determination of the frequency of spontaneous suppressor emergence showed a 13 to 25 times lower frequency for *tif51A-1* and *tif51A-3* mutants without than with the second eIF5A gene (*TIF51B*), respectively (Fig. 7a and Fig. 1a).

**Figure 7.**
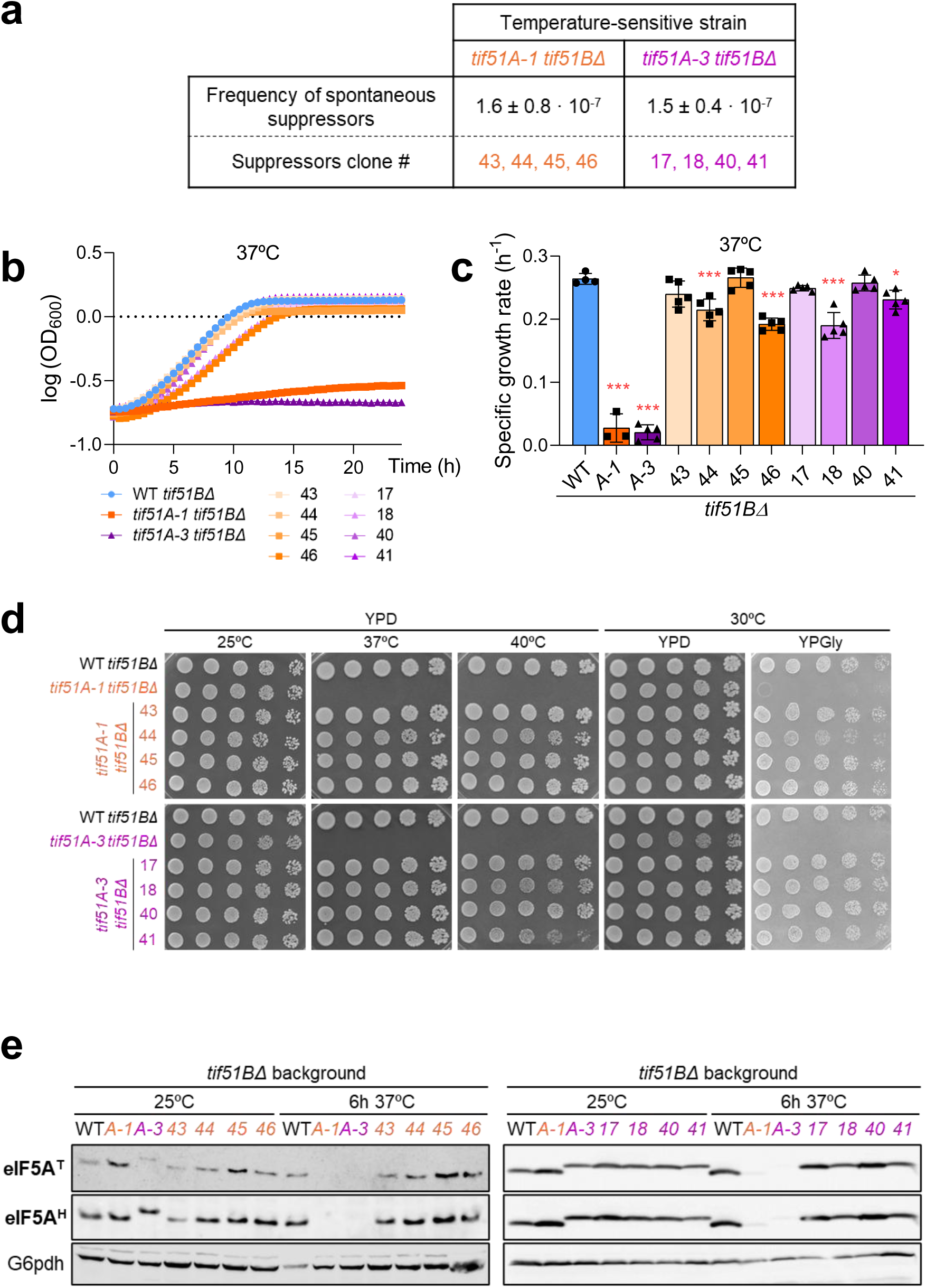
Isolation of suppressors of temperature-sensitive mutations in the yeast eIF5A encoding gene *TIF51A* in strains with deletion of *TIF51B* gene. (a) Frequency of spontaneous suppressor emergence. Frequencies were determined by growing *tif51A-1 tif51BΔ* and *tif51A-3 tif51BΔ* yeast cells in YPD to mid-exponential phase at 25°C and plating onto YPD agar. Plates were incubated at 25°C (permissive) and 37°C (restrictive) for 2 days. Frequencies were calculated as the ratio of colonies growing at 37°C relative to the total number of viable cells at 25°C. Specific independent suppressor clones selected for further characterization are identified by the numbers in orange (*tif51A-1 tif51BΔ* background) and purple (*tif51A-3 tif51BΔ* background). **(b, c)** Growth kinetics of suppressor strains at restrictive temperature. Wild-type (WT) *tif51BΔ*, *tif51A-1 tif51BΔ* and *tif51A-3 tif51BΔ* strains, along with their respective suppressors, were cultured in YPD at 25°C. Cultures were then diluted to an OD_600_ of 0.1 and shifted to 37°C. Growth was monitored by recording OD_600_ every 30 min for 24 h using a SPECTROstar Nano plate reader. Growth curves (b) are represented as the mean of log_10_ (OD_600_) from at least three biological replicates, and specific growth rates (c) were calculated from the slope of the exponential phase. Statistical significance was determined using a one-way ANOVA relative to the wild-type strain followed by Bonferroni’s post hoc multiple-comparison test. *p < 0.05, **p < 0.01, ***p < 0.001. Growth curves with error bars are provided in supplementary Figure S2. **(d)** Spot assay of suppressor strains. Same yeast strains used in (b, c) were cultured in YPD at 25°C and 5-fold serial dilutions were spotted onto YPD and YPGly plates. Plates were incubated for 2 – 3 days at 25°C, 37°C, and 40°C to assess temperature sensitivity, and at 30°C on YPGly to evaluate respiratory capacity. **(e)** Same yeast strains used in (b, c) were cultured in YPD at 25°C to mid-exponential phase at 25°C, and then shifted to 37°C for 6 h. Tif51A protein levels were determined by Western blotting using an anti-eIF5A antibody (which detects total Tif51A; eIF5A^T^), and an anti-hypusine antibody (which detects hypusinated Tif51A; eIF5A^H^). G6pdh protein levels were used as a loading control.

For the initial characterization of the suppressors, we studied their growth in glucose rich media at permissive (25°C), restrictive (37°C) and severe (40°C) temperatures, and also their ability to grow in a respiratory obligated media with glycerol as carbon source. All suppressors, but not the original *TIF51A* mutant strains, were able to grow under all conditions tested, and showed a similar or slightly reduced grow rate at restrictive temperature compared to the wild-type (Fig. 7b, c and d; and Supplementary Fig. S2). Furthermore, we examined the levels of the Tif51A protein in the different strains. After incubating the cells for 6 h at 37°C, Tif51A was undetectable in the parental mutant *tif51A-1* and *tif51A-3* cells, whereas high levels were observed in the wild-type cells. Unexpectedly, all suppressor cells exhibited Tif51A levels comparable to those of the wild-type, suggesting that the protein is stable at restrictive temperature and implying that the growth of the suppressors is attributable to normal Tif51A levels (Figure 7e).

To understand that result, we investigated the stability of the Tif51A protein in the wild-type, *tif51A-1* and *tif51A-3* mutant parental strains, and the suppressor strains, at restrictive temperature after translation shutdown with cycloheximide (CHX). The levels of native (wild-type, WT) Tif51A protein remained stable without any apparent reduction after 3 h of CHX treatment, whereas, as expected, Tif51A-1 and Tif51A-3 mutant proteins were undetectable after 1.5-2.0 h incubation at 37°C under CHX treatment. However, we found an important increase in Tif51A protein stability for all suppressors with differences between them (Fig. 8a). To investigate whether these changes in Tif51A protein stability were due to reversion of the mutations in the parental strains, or by the contrary were due to new mutations in other genes/pathways that affect the stability of Tif51A, we PCR amplified and sequenced the *TIF51A* gene for all suppressors. For the suppressors of the *tif51A-1 tif51BΔ* background, we found two of them (sups. 43 and 44) in which the point mutation of the parental strain Tif51A-1 (Pro83 to Ser) was reversed (Ser83 to Pro) (Fig. 8a, second upper panel); whereas in the other two (sups. 45 and 46) the original mutation (Pro83 to Ser) was present but a new mutation (Asp72 to Tyr) appeared (Fig. 8a, third upper panel). For the suppressors of the *tif51A-3 tif51BΔ* background, we found three of them in which one of the two-point mutations of the parental strain Tif51A-3 (Cys39 to Tyr) was reversed (Tyr39 to Cys; sups. 17 and 40) or mutation produced a change to a different amino acid (Tyr39 to Ser; sup. 18) (Fig. 8a, fifth and sixth panels from the top); whereas in the last suppressor (sup. 41), Tif51A protein kept the two parental mutations (Cys39 to Tyr and Gly118 to Asp) but the protein contained an additional mutation (Asp72 to Tyr) (Fig. 8a, lower panel). Interestingly, this mutation affecting the 72 amino acid residue was found in suppressors of both, *tif51A-1* and *tif51A-3* temperature-sensitivity mutations.

**Figure 8.**
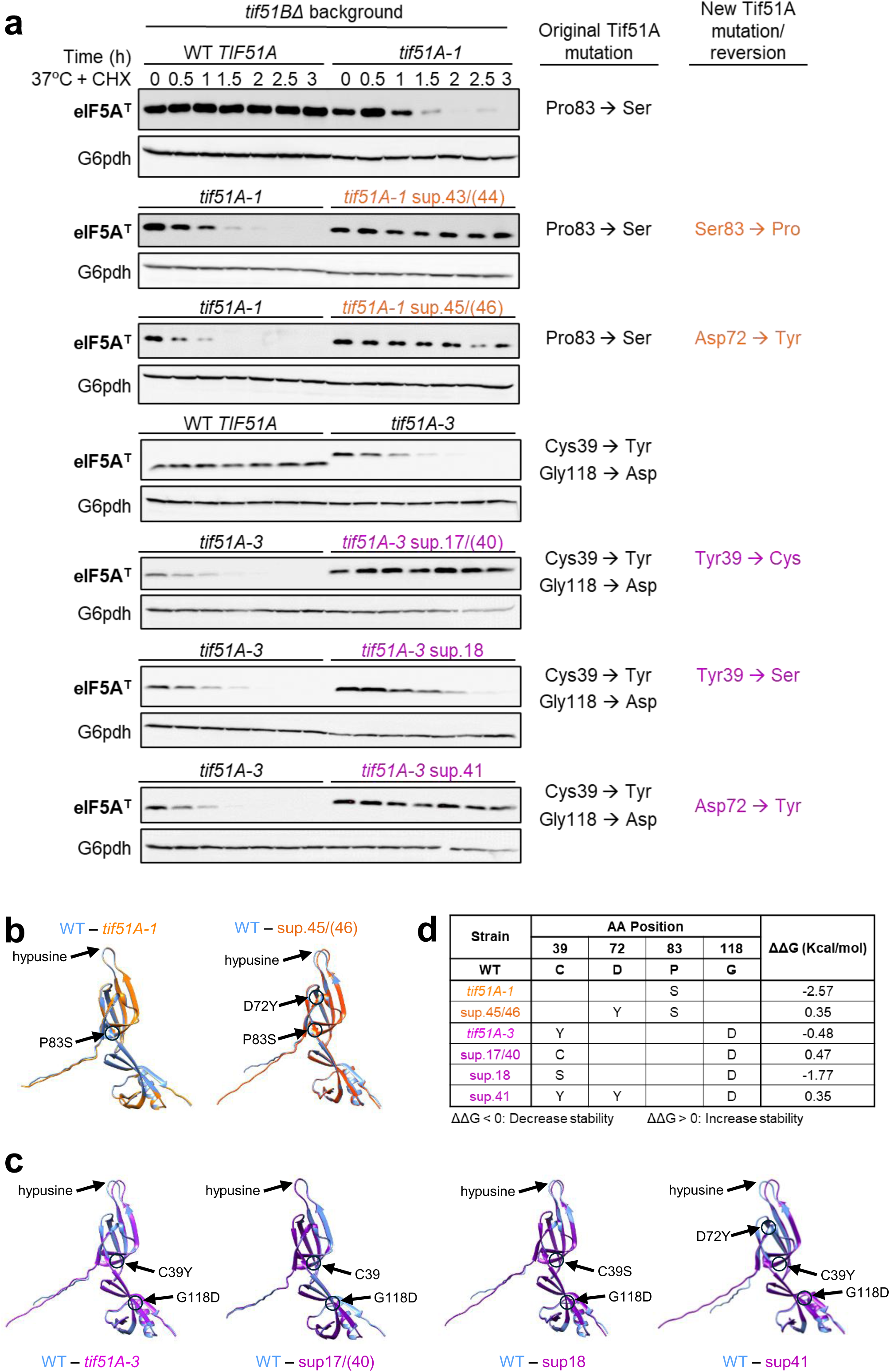
Suppressors of *tif51A-1/-3* mutations in strains with *tif51BΔ* background contain reversions or new mutations of *TIF51A* gene. (a) Wild-type (WT), *tif51A-1*, *tif51A-3*, and their respective suppressors were cultured in YPD to mid-exponential phase at 25°C, then shifted to 37°C for 3 h in the presence of 100 µg/mL CHX to inhibit *de novo* protein synthesis. Total eIF5A protein levels were determined by Western blotting using an anti-eIF5A antibody. G6pdh protein levels were used as a loading control. A representative image of one of the suppressors is shown. Sanger sequencing of the *TIF51A* ORF identified specific amino acid substitutions corresponding to either reversions or second-site mutations, correlating with the restoration of Tif51A protein stability under restrictive conditions. **(b, c)** AlphaFold2-predicted structural superpositions of Tif51A variants showing the alignment of the WT reference (blue) with *tif51A-1* derivatives (b) and *tif51A-3* derivatives (c). **(d)** Summary of the identified mutations and their predicted changes in folding free energy (ΔΔG, kcal/mol). Negative values (ΔΔG < 0) indicate decreased protein stability, while positive values (ΔΔG > 0) indicate increased protein stability relative to the WT.

To gain insight into the effects of mutations in the eIF5A protein tridimensional structure in the different Tif51A protein versions, we predicted the 3D-protein structure using AlphaFold2, and then, we visualized and compared the structures with UCSF Chimera. To estimate changes in protein stability, we calculated the folding free energy (ΔΔG, kcal/mol) in Biofold (see Materials and Methods for more information). We observed that Tif51A-1 and the native protein overlapped completely (Fig. 8b); whereas Tif51A-3 protein did not completely overlap with the native version, specifically in the area of the hypusination loop (Fig. 8c). For both Tif51A-1/-3 mutant proteins, the folding free energy was lower than that of the wild-type, therefore predicting decreased protein stability (Fig. 8d). These results correlate with the temperature-sensitive phenotypes of both Tif51A mutant proteins (Fig. 8a) and may also explain the more severe defects in growth of the *tif51A-3* mutant that it is observed also under permissive temperature (see in (Li et al. 2014; Barba-Aliaga et al. 2020)) that could be due to a partial loss of functionality of Tif51A carrying the two-point mutations (C39Y and G118D). For suppressors 17/40 for which only one of the two-point mutations of *tif51A-3* was reverted, the predicted protein structure shows a good overlapping with the native protein, pointing to the mutation in Cys39 as the one causing the more deleterious effect in protein functionality and stability (Fig. 8c). For the suppressors in which new mutations in Tif51A protein were identified we observed different results. For suppressors 45/46 of Tif51A-1 with new Asp72 to Tyr amino acid change, the predicted protein structure did not completely overlap with the native wild-type protein structure (Fig. 8b); however, there is a recovery in protein stability as estimated by the increase in the ΔΔG value (Fig. 8d), what correlates with the observed increase in protein stability (Fig. 8a). Similar results were observed for suppressor 41 of *tif51A-3*, which carries the same new mutation resulting in the amino acid change Asp72 to Tyr: the protein structure did not overlap completely with the native wild-type protein, but the predicted increase in stability correlated with the experimental data (Fig. 8c and d). Lastly, for suppressor 18 of the *tif51A-3* mutant, in which the first point mutation resulted in a changed to a different amino acid (Cys39 to Ser), a protein structure that overlaps with the native Tif51A protein was predicted; however, the ΔΔG value indicated low protein stability, which correlated with the partial restoration of protein stability (Fig. 8a, c and d). According to these data, suppressor 18 did not fully restore growth at 37°C either (see Fig. 7b and c; and Supplementary Fig. S2).

In sum, we can conclude that in cells with mutations in the *TIF51A* gene and deletion of the second eIF5A encoding gene, *TIF51B*, the appearance of suppressor showed a low frequency rate (of around 10^-7^) and no loss-of-function mutations were isolated that could suppress the lack of growth at 37°C. However, the frequency rate was compatible with reversion or gain-of-function mutations. Indeed, all isolated suppressors contained either reversions or compensatory mutations in the *TIF51A* coding sequence, which restored total or partially Tif51A protein stability and allowed for growth at 37°C.

## DISCUSSION

Over the past few decades, eIF5A has attracted significant attention due to its association with various diseases and conditions, including diabetes, cancer, neurodevelopmental disorders, viral infections and aging (Mathews and Hershey 2015; Ning et al. 2020; Tauc et al. 2021; Nakanishi and Cleveland 2024). eIF5A has been conserved throughout evolution and, in most eukaryotes, including yeast and humans, it is encoded by two paralogous genes as a result of ancient genome duplication events (Kang et al. 1992; Jenkins et al. 2001). Whilst most genes with paralogue copy are not deemed essential due to the secondary copy’s ability to serve as a compensatory backup mechanism (Seoighe and Wolfe 1999; Gu et al. 2003; Ihmels et al. 2007; Stein and Aloy 2008), the *TIF51A*/*EIF5A1* copy, which is highly expressed under normal aerobic conditions in yeast and most human tissues, respectively, is considered essential (Jenkins et al. 2001; Clement et al. 2003). Its unique characteristics also include the fact that it is the only know protein modified by hypusination, and yet its hypusination enzymes are also highly conserved and essential in eukaryotes (Park and Wolff 2018; Park et al. 2022). However, the essential function of eIF5A is not clearly assigned, although it is widely assumed that its main role is to function as a translation elongation factor, promoting the synthesis of hard-to translate amino acids (Dever et al. 2014; Dever et al. 2018; Park and Wolff 2018; Barba-Aliaga and Alepuz 2022a; Park et al. 2022). In this study, we aimed to use a genetic screening in yeast cells to identify genes and pathways that could compensate for the absence of a functional eIF5A protein in the cells, thereby providing more insight into the essential function of this factor. However, screening for suppressors of temperature-sensitive *TIF51A* mutations revealed that all the isolated suppressors, with frequency of appearance corresponding to loss-of-function mutations, contained mutations in the Rox1 and Mot3 repressor proteins, resulting in the up-regulation of the second eIF5A copy gene, *TIF51B*. We then performed a second screening using temperature-sensitive mutations in *TIF51A* in cells devoid of *TIF51B*. However, all isolated suppressors, with frequency of appearance corresponding to gain-of-function mutations, contained reversions of the *TIF51A* point mutations or new intragenic mutations that suppressed the temperature-dependent degradation of the Tif51A protein. Overall, the results demonstrate that there are no suppressors of eIF5A except eIF5A itself.

It is noteworthy that the results obtained indicated that the appearance of suppressors of Tif51A mutants was more prevalent in cells containing the second eIF5A gene, *TIF51B* (Fig. 1a). Consequently, novel point mutations (SNPs) or deletions and insertion (INDEL) mutations in the genes encoding the repressors Rox1 and Mot3 were observed to occur at a higher frequency than the reversion of mutations in *TIF51A*. These results suggest that the presence of the paralogue gene *TIF51B* acts as a backup mechanism to mitigate the fitness costs associated with the essential gene *TIF51A*. However, this compensatory effect does not occur within a cell via a direct feedback regulatory mechanism, as reported for other duplicated genes in yeast (Vande Zande et al. 2023) and plants (Nimchuk et al. 2015), but rather at the cell population level through the emergence of mutations that alter the regulation of the duplicated gene (Kafri et al. 2006).

It has been documented that the expression level of the Rox1 and Mot3 proteins is kept low in steady-state normoxic conditions, and that the two proteins must therefore act in synergy to achieve strong repression of *TIF51B* (Sertil 2003). In fact, Rox1 functions as a direct repressor of its own gene, thereby maintaining low levels of Rox1 protein and enabling rapid reprogramming of transcription in response to environmental changes (Deckert et al. 1995; Denby et al. 2012). Our findings indicate that the tight repression exerted by the joint action of Rox1 and Mot3, through their binding to multiple sites in the *TIF51B* promoter, allows both, to evade the deleterious effects associated with the co-expression of Tif51A and Tif51B in cells growing under aerobic conditions (Fig. 6); and to stablish a resettable repressor system over *TIF51B* that can be reversed by mutations in either Rox1 or Mot3.

The human *EIF5A2* gene, whose corresponding protein is 79.6% homologous to the yeast Tif51B protein, is repressed in most human tissues, but is expressed in the brain and testes. However, *EIF5A2* is up-regulated in many cancer types, where it has been proposed to promote proliferation, epithelial-mesenchymal transition (EMT), cellular migration, metastasis, and drug resistance. Therefore, despite the fact that EIF5A1 and EIF5A2 proteins share 94% similarity and overlapping functions, these paralogues also exhibit distinct roles in cellular processes and disease progression (Mathews and Hershey 2015; Ning et al. 2020; Nakanishi and Cleveland 2024; Xiong et al. 2025). In particular, EIF5A2 overexpression during human hepatocellular carcinoma (hHCC) development appears to drive metabolic reprogramming towards anaerobic glycolysis, with increased glucose uptake and lactate secretion, which are two metabolic hallmarks of cancer cells (Cao et al. 2017; Hanahan 2022). In contrast, human EIF5A1 promotes aerobic glycolysis with mitochondrial respiration (Puleston et al. 2019; Tauc et al. 2021). This functional distinction between the human eIF5A isoforms resembles the dichotomy of the yeast aerobic *TIF51A* and hypoxic *TIF51B* paralogues (Barba-Aliaga and Alepuz 2022a). Nevertheless, there is a scarcity of knowledge regarding the regulation of the *EIF5A2* gene. Notably, an orthologue of Rox1 has been identified in the human genome: the transcription factor SOX12 (Dy et al. 2008; Hoser et al. 2008). However, the involvement of SOX12 in regulating the human *EIF5A2* gene has not been described. Understanding the repressor mechanism acting on the *EIF5A2* gene in most tissues, and how this is cancelled during tumor progression may be crucial for elucidating therapeutic interventions in metastatic cancers.

The eIF5A isoform expressed under aerobic conditions in yeast, Tif51A, exhibits a high degree of conservation through evolution. Indeed, the *S. cerevisiae* protein shows 61.8% identity (82.8% similarity) with the human *EIF5A1* (Supplementary Fig. S3a and b). The Tif51A protein is highly stable (Fig. 8a), however, point mutations P83S (present in the *tif51A-1* mutant), and C39Y and G118D (both present in the *tif51A-3* mutant) result in loss of stability. These three amino acid residues are conserved from yeast to humans (Supplementary Fig. S3a), thus underscoring their significance for the structure and function of eIF5A. Residues C39 and P83 are located in two of the six β-strands (β3 and β6, respectively), which, together with the hypusination loop, constitute the N-terminal part of Tif51A protein; while residue G118 is located just before the only α-helix that forms part, along with five additional β-strands, the C-terminal part of Tif51A protein. In the present study, the suppressors of the *TIF51A* temperature-sensitive mutations isolated in *tif51BΔ* cells showed reversion of the mutation in P83 or C39, but not in G118, indicating that this last mutation alone does not induce Tif51A instability. Furthermore, suppressors of *tif51A-1* and *tif51A-3* were identified in which the original point mutations were present, but the appearance of the mutation D72Y, which is also found in an amino acid that is conserved from yeast to humans (located in the β5-strand, Supplementary Fig. S3a), restored protein stability (Fig. 8a). The amino acid sequence, 3D structure and hypusination of Tif51A, conserved in all eukaryotes, explain its ability to bind to the ribosome, locating next to the P-site (Melnikov et al. 2016; Schmidt et al. 2016). This high evolutionary conservation also explains why human, mouse and fly eIF5A can compensate for the absence of the protein in yeast cells (Schnier et al. 1991; Muñoz-Soriano et al. 2017). Although it is plausible that the essential role of eIF5A is linked to its interaction with the ribosome, there is no definitive explanation for its essentiality. However, our results show that, unlike other essential eukaryotic proteins, the eIF5A protein is indispensable. Therefore, our findings suggest that eIF5A performs an irreplaceable function within the ribosome, which cannot be substituted by loss- or gain-of-function mutations in any other protein in eukaryotic cells, including ribosomal components or any other translation factors.

## DATA AVAILABILITY

Strains and plasmids are available upon request. File S1 contains the supplementary tables (Tables S1-S5). File S2 contains the supplementary Materials and Methods, which provide detailed descriptions. File S3 contains supplementary Figures (Figures S1-S3). Whole Genome Sequencing (WGS) datasets are deposited in NCBI BioSample database under the accession number PRJNA1418127. All custom scripts used for bioinformatic analyses are available at https://github.com/Alejandro-121/Scripts-asociated-to-Prieto-D-ez-et-al-2026.

## ACKNOWLEDGMENTS

We thank Elena Valera (IATA-CSIC, València, Spain) for technical assistance with the CRISPR-Cas9 tools. We thank Ana María Pérez Adrián for technical assistance during genomic DNA extractions. We than Sebastián Chávez, Marina Barba-Aliaga and José Enrique Pérez-Ortín for critical reading of the manuscript. Whole Genome sequencing service was provided by Foundation for the Promotion of Health and Biomedical Research of the Valencian Community (FISABIO). We thank the Scientific Computing Area of Secretaría General Adjunta de Informática (SGAI)-CSIC for providing the computational resources used in this work (HPC-Drago high-performance computing cluster).

## CONFLICT OF INTEREST

The authors declare that they have followed the uniform requirements for manuscripts submitted to Biomedical journals. The authors declare no conflict of interest.

### Author Contributions

P.A. and S.P-D designed research; S.P-D, L.A. performed research; S.P-D., A.A., D.P. and P.A. analyzed data; P.A. wrote the paper. All authors revised the paper.

---

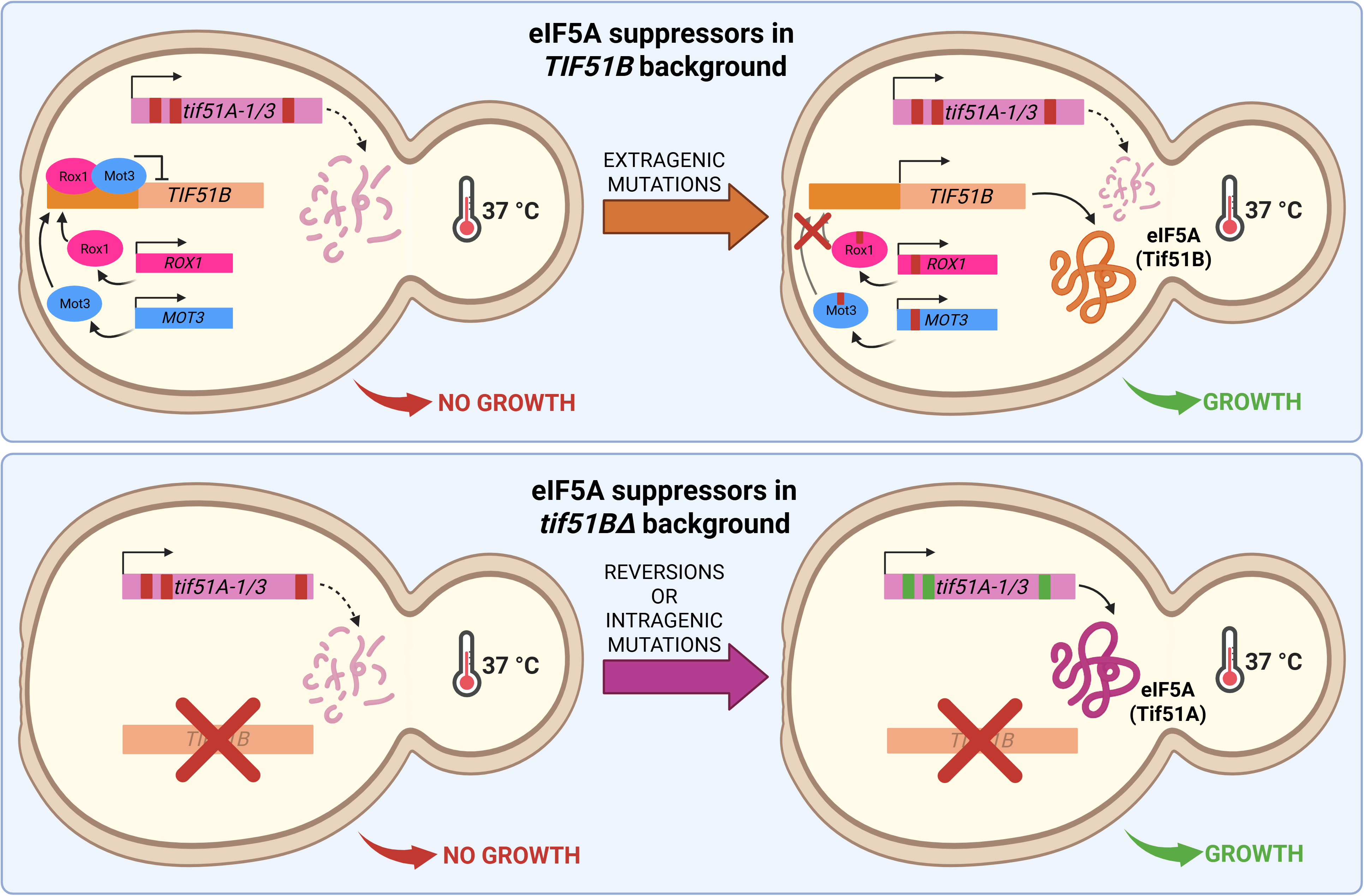

## REFERENCES

Abramova NE, Cohen BD, Sertil O, Kapoor R, Davies KJA, Lowry CV. 2001. Regulatory Mechanisms Controlling Expression of the *DAN* / *TIR* Mannoprotein Genes During Anaerobic Remodeling of the Cell Wall in *Saccharomyces cerevisiae*. Genetics. 157(3):1169–1177. 10.1093/genetics/157.3.1169

Aksu M, Trakhanov S, Vera Rodriguez A, Görlich D. 2019. Structural basis for the nuclear import and export functions of the biportin Pdr6/Kap122. J Cell Biol. 218(6):1839–1852. 10.1083/jcb.201812093

Alimardani P, Régnacq M, Moreau-Vauzelle C, Ferreira T, Rossignol T, Blondin B, Bergès T. 2004. *SUT1*-promoted sterol uptake involves the ABC transporter Aus1 and the mannoprotein Dan1 whose synergistic action is sufficient for this process. Biochem J. 381(1):195–202. 10.1042/BJ20040297

Auwera GV der, O’Connor BD. 2020. Genomics in the cloud: using Docker, GATK, and WDL in Terra. First edition. O’Reilly.

Barba-Aliaga M, Alepuz P. 2022a. Role of eIF5A in Mitochondrial Function. Int J Mol Sci. 23(3):1284. 10.3390/ijms23031284

Barba-Aliaga M, Alepuz P. 2022b. The activator/repressor Hap1 binds to the yeast eIF5A-encoding gene *TIF51A* to adapt its expression to the mitochondrial functional status. FEBS Lett. 596(14):1809–1826. 10.1002/1873-3468.14366

Barba-Aliaga M, Bernal V, Rong C, Volfbeyn ME, Zhang K, Zid BM, Alepuz P. 2024. eIF5A controls mitoprotein import by relieving ribosome stalling at *TIM50* translocase mRNA. J Cell Biol. 223(12):e202404094. 10.1083/jcb.202404094

Barba-Aliaga M, Chi L, Prieto-Díez S, Planells J, García-Martínez J, Pérez-Ortín JE, Alepuz P. 2026. eIF5A coordinates the transcription and translation of its target genes. Cell Mol Life Sci. Epub ahead of print. 10.1007/s00018-026-06252-8

Barba-Aliaga M, Mena A, Espinoza V, Apostolova N, Costell M, Alepuz P. 2021. Hypusinated eIF5A is required for the translation of collagen. J Cell Sci. 134(18):jcs258643. 10.1242/jcs.258643

Barba-Aliaga M, Villarroel-Vicente C, Stanciu A, Corman A, Martínez-Pastor MT, Alepuz P. 2020. Yeast Translation Elongation Factor eIF5A Expression Is Regulated by Nutrient Availability through Different Signalling Pathways. Int J Mol Sci. 22(1):219. 10.3390/ijms22010219

Bolotin-Fukuhara M, Casaregola S, Aigle M. 2005. Genome evolution: Lessons from Genolevures. In: Sunnerhagen P, Piskur J, (editors) Comparative Genomics. Topics in Current Genetics. Vol. 15. Springer Berlin Heidelberg; p 165–196. 10.1007/b136677

Burhans DT, Ramachandran L, Wang J, Liang P, Patterton HG, Breitenbach M, Burhans WC. 2006. Non-random clustering of stress-related genes during evolution of the *S. cerevisiae* genome. BMC Evol Biol. 6(1):58. 10.1186/1471-2148-6-58

Cao T-T, Lin S-H, Fu L, Tang Z, Che C-M, Zhang L-Y, Ming X-Y, Liu T-F, Tang X-M, Tan B-B, et al. 2017. Eukaryotic translation initiation factor 5A2 promotes metabolic reprogramming in hepatocellular carcinoma cells. Carcinogenesis. 38(1):94–104. 10.1093/carcin/bgw119

Capriotti E, Fariselli P, Casadio R. 2005. I-Mutant2.0: predicting stability changes upon mutation from the protein sequence or structure. Nucleic Acids Res. 33(suppl_2):W306–W310. 10.1093/nar/gki375

Caraglia M, Park MH, Wolff EC, Marra M, Abbruzzese A. 2013. eIF5A isoforms and cancer: two brothers for two functions? Amino Acids. 44(1):103–109. 10.1007/s00726-011-1182-x

Cherry J. 1998. SGD: Saccharomyces Genome Database. Nucleic Acids Res. 26(1):73–79. 10.1093/nar/26.1.73

Choi Y, Um B, Na Y, Kim J, Kim J-S, Kim VN. 2024. Time-resolved profiling of RNA binding proteins throughout the mRNA life cycle. Mol Cell. 84(9):1764–1782.e10. 10.1016/j.molcel.2024.03.012

Cingolani P, Platts A, Wang LL, Coon M, Nguyen T, Wang L, Land SJ, Lu X, Ruden DM. 2012. A program for annotating and predicting the effects of single nucleotide polymorphisms, SnpEff: SNPs in the genome of Drosophila melanogaster strain w^1118^; iso-2; iso-3. Fly. 6(2):80–92. 10.4161/fly.19695

Clement PMJ, Henderson CA, Jenkins ZA, Smit-McBride Z, Wolff EC, Hershey JWB, Park MH, Johansson HE. 2003. Identification and characterization of eukaryotic initiation factor 5A-2. Eur J Biochem. 270(21):4254–4263. 10.1046/j.1432-1033.2003.03806.x

Deckert J, Rodriguez Torres AM, Simon JT, Zitomer RS. 1995. Mutational Analysis of Rox1, a DNA-Bending Repressor of Hypoxic Genes in *Saccharomyces cerevisiae*. Mol Cell Biol. 15(11):6109–6117. 10.1128/MCB.15.11.6109

Denby CM, Im JH, Yu RC, Pesce CG, Brem RB. 2012. Negative feedback confers mutational robustness in yeast transcription factor regulation. Proc Natl Acad Sci. 109(10):3874–3878. 10.1073/pnas.1116360109

Dever TE, Dinman JD, Green R. 2018. Translation Elongation and Recoding in Eukaryotes. Cold Spring Harb Perspect Biol. 10(8):a032649. 10.1101/cshperspect.a032649

Dever TE, Gutierrez E, Shin B-S. 2014. The hypusine-containing translation factor eIF5A. Crit Rev Biochem Mol Biol. 49(5):413–425. 10.3109/10409238.2014.939608

Dy P, Penzo-Méndez A, Wang H, Pedraza CE, Macklin WB, Lefebvre V. 2008. The three SoxC proteins--Sox4, Sox11 and Sox12--exhibit overlapping expression patterns and molecular properties. Nucleic Acids Res. 36(9):3101–3117. 10.1093/nar/gkn162

Fang Z-A, Wang G-H, Chen A-L, Li Y-F, Liu J-P, Li Y-Y, Bolotin-Fukuhara M, Bao W-G. 2009. Gene Responses to Oxygen Availability in *Kluyveromyces lactis*: an Insight on the Evolution of the Oxygen-Responding System in Yeast. PLoS ONE. 4(10):e7561. 10.1371/journal.pone.0007561

Garre E, Romero-Santacreu L, Barneo-Muñoz M, Miguel A, Pérez-Ortín JE, Alepuz P. 2013. Nonsense-Mediated mRNA Decay Controls the Changes in Yeast Ribosomal Protein Pre-mRNAs Levels upon Osmotic Stress. PLoS ONE. 8(4):e61240. 10.1371/journal.pone.0061240

Grishin AV, Rothenberg M, Downs MA, Blumer KJ. 1998. Mot3, a Zn Finger Transcription Factor That Modulates Gene Expression and Attenuates Mating Pheromone Signaling in *Saccharomyces cerevisiae*. Genetics. 149(2):879–892. 10.1093/genetics/149.2.879

Gu Z, Steinmetz LM, Gu X, Scharfe C, Davis RW, Li W-H. 2003. Role of duplicate genes in genetic robustness against null mutations. Nature. 421(6918):63–66. 10.1038/nature01198

Gutierrez E, Shin B-S, Woolstenhulme CJ, Kim J-R, Saini P, Buskirk AR, Dever TE. 2013. eIF5A Promotes Translation of Polyproline Motifs. Mol Cell. 51(1):35–45. 10.1016/j.molcel.2013.04.021

Hanahan D. 2022. Hallmarks of Cancer: New Dimensions. Cancer Discov. 12(1):31–46. 10.1158/2159-8290.CD-21-1059

Hickman MJ, Winston F. 2007. Heme Levels Switch the Function of Hap1 of *Saccharomyces cerevisiae* between Transcriptional Activator and Transcriptional Repressor. Mol Cell Biol. 27(21):7414–7424. 10.1128/MCB.00887-07

Hittinger CT, Carroll SB. 2007. Gene duplication and the adaptive evolution of a classic genetic switch. Nature. 449(7163):677–681. 10.1038/nature06151

Hodge MR, Kim G, Singh K, Cumsky MG. 1989. Inverse regulation of the yeast *COX5* genes by oxygen and heme. Mol Cell Biol. 9(5):1958–1964. 10.1128/MCB.9.5.1958

Hoser M, Potzner MR, Koch JMC, Bösl MR, Wegner M, Sock E. 2008. Sox12 Deletion in the Mouse Reveals Nonreciprocal Redundancy with the Related Sox4 and Sox11 Transcription Factors. Mol Cell Biol. 28(15):4675–4687. 10.1128/MCB.00338-08

Ihmels J, Collins SR, Schuldiner M, Krogan NJ, Weissman JS. 2007. Backup without redundancy: genetic interactions reveal the cost of duplicate gene loss. Mol Syst Biol. 3(1):86. 10.1038/msb4100127

Ishfaq M, Maeta K, Maeda S, Natsume T, Ito A, Yoshida M. 2012. Acetylation regulates subcellular localization of eukaryotic translation initiation factor 5A (eIF5A). FEBS Lett. 586(19):3236–3241. 10.1016/j.febslet.2012.06.042

Jenkins ZA, Hååg PG, Johansson HE. 2001. Human EIF5A2 on Chromosome 3q25–q27 Is a Phylogenetically Conserved Vertebrate Variant of Eukaryotic Translation Initiation Factor 5A with Tissue-Specific Expression. Genomics. 71(1):101–109. 10.1006/geno.2000.6418

Kafri R, Levy M, Pilpel Y. 2006. The regulatory utilization of genetic redundancy through responsive backup circuits. Proc Natl Acad Sci. 103(31):11653–11658. 10.1073/pnas.0604883103

Kang HA, Schwelberger HG, Hershey JWB. 1992. The two genes encoding protein synthesis initiation factor eIF-5A in *Saccharomyces cerevisiae* are members of a duplicated gene cluster. Mol Gen Genet MGG. 233(3):487–490. 10.1007/BF00265449

Kastaniotis AJ, Mennella TA, Konrad C, Torres AMR, Zitomer RS. 2000. Roles of Transcription Factor Mot3 and Chromatin in Repression of the Hypoxic Gene *ANB1* in Yeast. Mol Cell Biol. 20(19):7088–7098. 10.1128/MCB.20.19.7088-7098.2000

Kastaniotis AJ, Zitomer RS. 2000. Rox1 mediated repression. Oxygen dependent repression in yeast. Adv Exp Med Biol. 475:185–195. 10.1007/0-306-46825-5_18

Kellis M, Birren BW, Lander ES. 2004. Proof and evolutionary analysis of ancient genome duplication in the yeast *Saccharomyces cerevisiae*. Nature. 428(6983):617–624. 10.1038/nature02424

Klinkenberg LG, Mennella TA, Luetkenhaus K, Zitomer RS. 2005. Combinatorial Repression of the Hypoxic Genes of *Saccharomyces cerevisiae* by DNA Binding Proteins Rox1 and Mot3. Eukaryot Cell. 4(4):649–660. 10.1128/EC.4.4.649-660.2005

Kowalski LRZ, Kondo K, Inouye M. 1995. Cold-shock induction of a family of TIP1-related proteins associated with the membrane in *Saccharomyces cerevisiae*. Mol Microbiol. 15(2):341–353. 10.1111/j.1365-2958.1995.tb02248.x

Kwast KE, Lai L-C, Menda N, James DT, Aref S, Burke PV. 2002. Genomic Analyses of Anaerobically Induced Genes in *Saccharomyces cerevisiae*: Functional Roles of Rox1 and Other Factors in Mediating the Anoxic Response. J Bacteriol. 184(1):250–265. 10.1128/JB.184.1.250-265.2002

Lai L-C, Kosorukoff AL, Burke PV, Kwast KE. 2006. Metabolic-State-Dependent Remodeling of the Transcriptome in Response to Anoxia and Subsequent Reoxygenation in *Saccharomyces cerevisiae*. Eukaryot Cell. 5(9):1468–1489. 10.1128/EC.00107-06

Langdon QK, Peris D, Kyle B, Hittinger CT. 2018. sppIDer: A Species Identification Tool to Investigate Hybrid Genomes with High-Throughput Sequencing. Mol Biol Evol. 10.1093/molbev/msy166

Levasseur EM, Yamada K, Piñeros AR, Wu W, Syed F, Orr KS, Anderson-Baucum E, Mastracci TL, Maier B, Mosley AL, et al. 2019. Hypusine biosynthesis in β cells links polyamine metabolism to facultative cellular proliferation to maintain glucose homeostasis. Sci Signal. 12(610):eaax0715. 10.1126/scisignal.aax0715

Li H. 2013. Aligning sequence reads, clone sequences and assembly contigs with BWA-MEM. 10.48550/arXiv.1303.3997

Li T, Belda-Palazón B, Ferrando A, Alepuz P. 2014. Fertility and Polarized Cell Growth Depends on eIF5A for Translation of Polyproline-Rich Formins in *Saccharomyces cerevisiae*. Genetics. 197(4):1191–1200. 10.1534/genetics.114.166926

Li T, De Clercq N, Medina DA, Garre E, Sunnerhagen P, Pérez-Ortín JE, Alepuz P. 2016. The mRNA cap-binding protein Cbc1 is required for high and timely expression of genes by promoting the accumulation of gene-specific activators at promoters. Biochim Biophys Acta BBA - Gene Regul Mech. 1859(2):405–419. 10.1016/j.bbagrm.2016.01.002

Liang Y, Piao C, Beuschel CB, Toppe D, Kollipara L, Bogdanow B, Maglione M, Lützkendorf J, See JCK, Huang S, et al. 2021. eIF5A hypusination, boosted by dietary spermidine, protects from premature brain aging and mitochondrial dysfunction. Cell Rep. 35(2):108941. 10.1016/j.celrep.2021.108941

Lowry CV, Cerdán ME, Zitomer RS. 1990. A hypoxic consensus operator and a constitutive activation region regulate the *ANB1* gene of *Saccharomyces cerevisiae*. Mol Cell Biol. 10(11):5921–5926. 10.1128/MCB.10.11.5921

Lowry CV, Zitomer RS. 1988. *ROX1* Encodes a Heme-Induced Repression Factor Regulating *ANB1* and *CYC7* of *Saccharomyces cerevisiae*. Mol Cell Biol. 8(11):4651–4658. 10.1128/mcb.8.11.4651-4658.1988

Lubas M, Harder LM, Kumsta C, Tiessen I, Hansen M, Andersen JS, Lund AH, Frankel LB. 2018. eIF5A is required for autophagy by mediating ATG3 translation. EMBO Rep. 19(6): e46072. 10.15252/embr.201846072

MacKintosh C, Ferrier DEK. 2018. Recent advances in understanding the roles of whole genome duplications in evolution. F1000Research. 6:1623. 10.12688/f1000research.11792.2

Madeira F, Madhusoodanan N, Lee J, Eusebi A, Niewielska A, Tivey ARN, Lopez R, Butcher S. 2024. The EMBL-EBI Job Dispatcher sequence analysis tools framework in 2024. Nucleic Acids Res. 52(W1):W521–W525. 10.1093/nar/gkae241

Maier B, Ogihara T, Trace AP, Tersey SA, Robbins RD, Chakrabarti SK, Nunemaker CS, Stull ND, Taylor CA, Thompson JE, et al. 2010. The unique hypusine modification of eIF5A promotes islet β cell inflammation and dysfunction in mice. J Clin Invest. 120(6):2156–2170. 10.1172/JCI38924

Marcet-Houben M, Gabaldón T. 2015. Beyond the Whole-Genome Duplication: Phylogenetic Evidence for an Ancient Interspecies Hybridization in the Baker’s Yeast Lineage. PLOS Biol. 13(8):e1002220. 10.1371/journal.pbio.1002220

Mathews MB, Hershey JWB. 2015. The translation factor eIF5A and human cancer. Biochim Biophys Acta BBA - Gene Regul Mech. 1849(7):836–844. 10.1016/j.bbagrm.2015.05.002

Melnikov S, Mailliot J, Shin B-S, Rigger L, Yusupova G, Micura R, Dever TE, Yusupov M. 2016. Crystal Structure of Hypusine-Containing Translation Factor eIF5A Bound to a Rotated Eukaryotic Ribosome. J Mol Biol. 428(18):3570–3576. 10.1016/j.jmb.2016.05.011

Mirdita M, Schütze K, Moriwaki Y, Heo L, Ovchinnikov S, Steinegger M. 2022. ColabFold: making protein folding accessible to all. Nat Methods. 19(6):679–682. 10.1038/s41592-022-01488-1

Muñoz-Soriano V, Domingo-Muelas A, Li T, Gamero E, Bizy A, Fariñas I, Alepuz P, Paricio N. 2017. Evolutionary conserved role of eukaryotic translation factor eIF5A in the regulation of actin-nucleating formins. Sci Rep. 7(1):9580. 10.1038/s41598-017-10057-y

Nakanishi S, Cleveland JL. 2024. The Many Faces of Hypusinated eIF5A: Cell Context-Specific Effects of the Hypusine Circuit and Implications for Human Health. Int J Mol Sci. 25(15):8171. 10.3390/ijms25158171

Nimchuk ZL, Zhou Y, Tarr PT, Peterson BA, Meyerowitz EM. 2015. Plant stem cell maintenance by transcriptional cross-regulation of related receptor kinases. Development. 142(6):1043–1049. 10.1242/dev.119677

Ning L, Wang L, Zhang H, Jiao X, Chen D. 2020. Eukaryotic translation initiation factor 5A in the pathogenesis of cancers (Review). Oncol Lett. 20(4):1–1. 10.3892/ol.2020.11942

Olsen ME, Connor JH. 2017. Hypusination of eIF5A as a Target for Antiviral Therapy. DNA Cell Biol. 36(3):198–201. 10.1089/dna.2016.3611

Park MH, Kar RK, Banka S, Ziegler A, Chung WK. 2022. Post-translational formation of hypusine in eIF5A: implications in human neurodevelopment. Amino Acids. 54(4):485–499. 10.1007/s00726-021-03023-6

Park MH, Wolff EC. 2018. Hypusine, a polyamine-derived amino acid critical for eukaryotic translation. J Biol Chem. 293(48):18710–18718. 10.1074/jbc.TM118.003341

Pelechano V, Alepuz P. 2017. eIF5A facilitates translation termination globally and promotes the elongation of many non polyproline-specific tripeptide sequences. Nucleic Acids Res. 45(12):7326–7338. 10.1093/nar/gkx479

Pettersen EF, Goddard TD, Huang CC, Couch GS, Greenblatt DM, Meng EC, Ferrin TE. 2004. UCSF Chimera—A visualization system for exploratory research and analysis. J Comput Chem. 25(13):1605–1612. 10.1002/jcc.20084

Puleston DJ, Buck MD, Klein Geltink RI, Kyle RL, Caputa G, O’Sullivan D, Cameron AM, Castoldi A, Musa Y, Kabat AM, et al. 2019. Polyamines and eIF5A Hypusination Modulate Mitochondrial Respiration and Macrophage Activation. Cell Metab. 30(2):352–363.e8. 10.1016/j.cmet.2019.05.003

Querol A, Barrio E, Ramón D. 1992. A Comparative Study of Different Methods of Yeast Strain Characterization. Syst Appl Microbiol. 15(3):439–446. 10.1016/S0723-2020(11)80219-5

Rosorius O, Reichart B, Krätzer F, Heger P, Dabauvalle M-C, Hauber J. 1999. Nuclear pore localization and nucleocytoplasmic transport of eIF-5A: evidence for direct interaction with the export receptor CRM1. J Cell Sci. 112(14):2369–2380. 10.1242/jcs.112.14.2369

Ruhl M, Himmelspach M, Bahr GM, Hammerschmid F, Jaksche H, Wolff B, Aschauer H, Farrington GK, Probst H, Bevec D. 1993. Eukaryotic initiation factor 5A is a cellular target of the human immunodeficiency virus type 1 Rev activation domain mediating trans-activation. J Cell Biol. 123(6):1309–1320. 10.1083/jcb.123.6.1309

Schmidt C, Becker T, Heuer A, Braunger K, Shanmuganathan V, Pech M, Berninghausen O, Wilson DN, Beckmann R. 2016. Structure of the hypusinylated eukaryotic translation factor eIF-5A bound to the ribosome. Nucleic Acids Res. 44(4):1944–1951. 10.1093/nar/gkv1517

Schnier J, Schwelberger HG, Smit-McBride Z, Kang HA, Hershey JW. 1991. Translation initiation factor 5A and its hypusine modification are essential for cell viability in the yeast *Saccharomyces cerevisiae*. Mol Cell Biol. 11(6):3105–3114. 10.1128/MCB.11.6.3105

Schuller AP, Wu CC-C, Dever TE, Buskirk AR, Green R. 2017. eIF5A Functions Globally in Translation Elongation and Termination. Mol Cell. 66(2):194–205.e5. 10.1016/j.molcel.2017.03.003

Schüller H-J. 2003. Transcriptional control of nonfermentative metabolism in the yeast *Saccharomyces cerevisiae*. Curr Genet. 43(3):139–160. 10.1007/s00294-003-0381-8

Schwelberger HG, Kang HA, Hershey JW. 1993. Translation initiation factor eIF-5A expressed from either of two yeast genes or from human cDNA. Functional identity under aerobic and anaerobic conditions. J Biol Chem. 268(19):14018–14025. 10.1016/S0021-9258(19)85203-1

Seoighe C, Wolfe KH. 1998. Extent of genomic rearrangement after genome duplication in yeast. Proc Natl Acad Sci. 95(8):4447–4452. 10.1073/pnas.95.8.4447

Seoighe C, Wolfe KH. 1999. Yeast genome evolution in the post-genome era. Curr Opin Microbiol. 2(5):548–554. 10.1016/S1369-5274(99)00015-6

Sertil O. 2003. Synergistic repression of anaerobic genes by Mot3 and Rox1 in *Saccharomyces cerevisiae*. Nucleic Acids Res. 31(20):5831–5837. 10.1093/nar/gkg792

Shaw W, Ellis T. 2018. Quick and easy CRISPR engineering in *Saccharomyces cerevisiae*. https://benchling.com/pub/ellis-crispr-tools#protocols

Stein A, Aloy P. 2008. A molecular interpretation of genetic interactions in yeast. FEBS Lett. 582(8):1245–1250. 10.1016/j.febslet.2008.02.020

Sullivan DP, Georgiev A, Menon AK. 2009. Tritium Suicide Selection Identifies Proteins Involved in the Uptake and Intracellular Transport of Sterols in *Saccharomyces cerevisiae*. Eukaryot Cell. 8(2):161–169. 10.1128/EC.00135-08

Tauc M, Cougnon M, Carcy R, Melis N, Hauet T, Pellerin L, Blondeau N, Pisani DF. 2021. The eukaryotic initiation factor 5A (eIF5A1), the molecule, mechanisms and recent insights into the pathophysiological roles. Cell Biosci. 11(1):219. 10.1186/s13578-021-00733-y

Tersey SA, Colvin SC, Maier B, Mirmira RG. 2014. Protective effects of polyamine depletion in mouse models of type 1 diabetes: implications for therapy. Amino Acids. 46(3):633–642. 10.1007/s00726-013-1560-7

Tesina P, Ebine S, Buschauer R, Thoms M, Matsuo Y, Inada T, Beckmann R. 2023. Molecular basis of eIF5A-dependent CAT tailing in eukaryotic ribosome-associated quality control. Mol Cell. 83(4):607–621.e4. 10.1016/j.molcel.2023.01.020

Trueblood CE, Poyton RO. 1988. Identification of *REO1*, a gene involved in negative regulation of *COX5b* and *ANB1* in aerobically grown *Saccharomyces cerevisiae*. Genetics. 120(3):671–680. 10.1093/genetics/120.3.671

Urban-Grimal D, Labbe-Bois R. 1981. Genetic and biochemical characterization of mutants of *Saccharomyces cerevisiae* blocked in six different steps of heme biosynthesis. Mol Gen Genet MGG. 183(1):85–92. 10.1007/BF00270144

Vande Zande P, Siddiq MA, Hodgins-Davis A, Kim L, Wittkopp PJ. 2023. Active compensation for changes in *TDH3* expression mediated by direct regulators of *TDH3* in *Saccharomyces cerevisiae*. PLOS Genet. 19(12):e1011078. 10.1371/journal.pgen.1011078

Wang F, Guan X, Xie D. 2013. Roles of Eukaryotic Initiation Factor 5A2 in Human Cancer. Int J Biol Sci. 9(10):1013–1020. 10.7150/ijbs.7191

Waterhouse AM, Procter JB, Martin DMA, Clamp M, Barton GJ. 2009. Jalview Version 2—a multiple sequence alignment editor and analysis workbench. Bioinformatics. 25(9):1189–1191. 10.1093/bioinformatics/btp033

Wilcox LJ, Balderes DA, Wharton B, Tinkelenberg AH, Rao G, Sturley SL. 2002. Transcriptional Profiling Identifies Two Members of the ATP-binding Cassette Transporter Superfamily Required for Sterol Uptake in Yeast. J Biol Chem. 277(36):32466–32472. 10.1074/jbc.M204707200

Wöhl T, Klier H, Ammer H, Lottspeich F, Magdolen V. 1993. The *HYP2* gene of *Saccharomyces cerevisiae* is essential for aerobic growth: characterization of different isoforms of the hypusine-containing protein Hyp2p and analysis of gene disruption mutants. Mol Gen Genet. 241(3–4):305–311. 10.1007/BF00284682

Wolfe KH, Shields DC. 1997. Molecular evidence for an ancient duplication of the entire yeast genome. Nature. 387(6634):708–713. 10.1038/42711

Xiong X, Du Y, Liu P, Li X, Lai X, Miao H, Ning B. 2025. Unveiling EIF5A2: A multifaceted player in cellular regulation, tumorigenesis and drug resistance. Eur J Pharmacol. 997:177596. 10.1016/j.ejphar.2025.177596

Xu A, Jao DL-E, Chen KY. 2004. Identification of mRNA that binds to eukaryotic initiation factor 5A by affinity co-purification and differential display. Biochem J. 384(3):585–590. 10.1042/BJ20041232

Zhang H, Alsaleh G, Feltham J, Sun Y, Napolitano G, Riffelmacher T, Charles P, Frau L, Hublitz P, Yu Z, et al. 2019. Polyamines Control eIF5A Hypusination, TFEB Translation, and Autophagy to Reverse B Cell Senescence. Mol Cell. 76(1):110–125.e9. 10.1016/j.molcel.2019.08.005

Zhang L, Hach A. 1999. Molecular mechanism of heme signaling in yeast: the transcriptional activator Hap1 serves as the key mediator. Cell Mol Life Sci. 56(5–6):415–426. 10.1007/s000180050442

Zitomer RS, Lowry CV. 1992. Regulation of gene expression by oxygen in *Saccharomyces cerevisiae*. Microbiol Rev. 56(1):1–11. 10.1128/mr.56.1.1-11.1992

Zuzuarregui A, Li T, Friedmann C, Ammerer G, Alepuz P. 2015. Msb2 is a Ste11 membrane concentrator required for full activation of the HOG pathway. Biochim Biophys Acta BBA - Gene Regul Mech. 1849(6):722–730. 10.1016/j.bbagrm.2015.02.001

